# Antimicrobial and Cytotoxic Lysolipins I–M Isolated from *Streptomyces* sp. P8-2B18

**DOI:** 10.64898/2026.06.12.731603

**Authors:** Manar Magdy Mahmoud Mahmoud, Kah Yean Lum, Yiwei Liu, Mikael Lenz Strube, Gundela Peschel, Luciano D. O. Souza, Miriam A. Rosenbaum, José M. A. Moreira, Stewart Brian Kirton, Charlotte H. Gotfredsen, Ling Ding

## Abstract

Lysolipin I (**1**) is a highly bioactive xanthone with strong antibacterial and cytotoxic properties. Given the limited number of lysolipin analogues, discovery of new natural lysolipin derivatives is important for understanding their structure-activity relationships. A soil-derived *Streptomyces* sp. P8-2B18 harbors a putative lysolipin biosynthetic gene cluster and LC-MS based metabolomic analysis revealed the production of lysolipin I along with unreported analogues. Large-scale fermentation followed by isolation led to the discovery of four new analogues, lysolipins J-M (**2**– **5**), the structures of which were elucidated by mass spectrometric and NMR spectroscopic data analyses. Lysolipin L features a five-membered lactam F ring, which was unprecedented in reported lysolipins. Lysolipin M has a novel skeleton, with an extra methyl (Me-36) and a glycosyl group replacing a 1,3-oxane ring in lysolipin I. While lysolipins I, J and K displayed strong activity against *Staphylococcus aureus* and *Aspergillus flavus* with MIC values ranging from 0.25 to 4 μg/mL and lysolipin L showed only moderate activities, lysolipin M was inactive (>50 μg/mL). Lysolipins I–K showed potent cytotoxic activity against prostate cancer cell lines LNCaP and C4-2B, with IC_50_ values in the submicromolar range. In contrast, lysolipin L exhibited no cytotoxicity and lysolipin M exhibited substantially reduced potency. Their broad, non-selective bioactivities restricted their applicability as therapeutic agents.

---

Polycyclic xanthone natural products constitute a class of polyketides characterized by their highly oxygenated angular hexacyclic ring systems. They have drawn interest owing to their unique chemical architecture and diverse biological activities. Structural diversity of the polycyclic xanthones arises from oxidation states of the xanthone ring, and its multifaceted substitutions including hydroxylation, halogenation and glycosylation, together with the presence of a methylene dioxybridge fused to the framework, and the quinone/hydroquinone oxidation state.^1^ This class of molecules have been isolated from various actinomycetes^2^ and the representative bioactive xanthones include lysolipins,^3^ cervinomycins,^4^ citreamicins,^5^ and actinoplanones.^6^

Lysolipins are chlorinated xanthones that were firstly isolated from *Streptomyces violaceoniger* Tü 96 in 1975.^3^ They are biosynthesized by a type 2 polyketide synthase (PKS), involving multiple cyclization steps of a polyketide precursor that give rise to their highly oxygenated aromatic and heteroaromatic rings.^7,8^ Extensive post-PKS oxidative modification further contributes to their structural complexity, as supported by its chemical structure and isotope feeding studies.^7^ To date, lysolipins I and X are the only naturally occurring members^3^ while several synthetic analogues have been patented,^9^ and new analogues generated by genetic engineering.^10^ Lysolipin I has demonstrated strong activity against Gram-positive and some Gram-negative bacteria. Studies suggested its role in interacting with the carrier lipid resulting in inhibition of cell wall biosynthesis, concluded by accumulation of murein precursors.^3^

Our recent investigation of *Streptomyces* derived from soil collected in the Jægersborg Deer Park, Denmark, resulted in discovery of several new specialized metabolites. For example, *Streptomyces mirabilis* strain P8-A2 produced azodyrecins A–C,^11^ pepticinnamins N–P,^12^ and maramycin.^13^ These findings motivated further exploration of other *Streptomyces* spp. derived from this environment. Through cocultivation, *Streptomyces* sp. P8-2B18 exhibited potent antifungal activities against several fungal pathogens and therefore was prioritized for further chemical investigations. An integrated approach combining genome-mining, metabolomic analysis, and bioassay-guided isolation resulted in discovery of several unprecedented lysolipin analogues. We hereby describe the isolation, structural identification, biological activities, and molecular docking studies of the isolated compounds.

## RESULTS AND DISCUSSION

Cocultivation revealed that a soil derived *Streptomyces* sp. P8-2B18 possessed antagonistic activity against several fungal pathogens, including *Fusarium graminearum, Alternaria solani, Aspergillus flavus, Aspergillus niger*, and *Aspergillus fumigatus* on PDA plates (**Figure S1**). Following HPLC-HRMS and Global Natural Product Social Molecular Networking (GNPS) metabolomic analysis, the EtOAc extract of the strain indicated the presence of lysolipin I as the major component, together with several unknown analogues (**Figure S2**). Based on initial screening using OSMAC (One Strain MAny Compounds) approach,^14^ the strain was fermented at a large scale (50 L) in ISP2 medium and the culture broth was fractionated through flash column chromatography on Amberchrom 161CGM chromatography resin. The fraction containing lysolipins was subjected to silica gel and Sephadex LH-20 column chromatography, followed by reversed-phase HPLC to yield compounds **1–5** (**Figure 1**).

**Figure 1.**
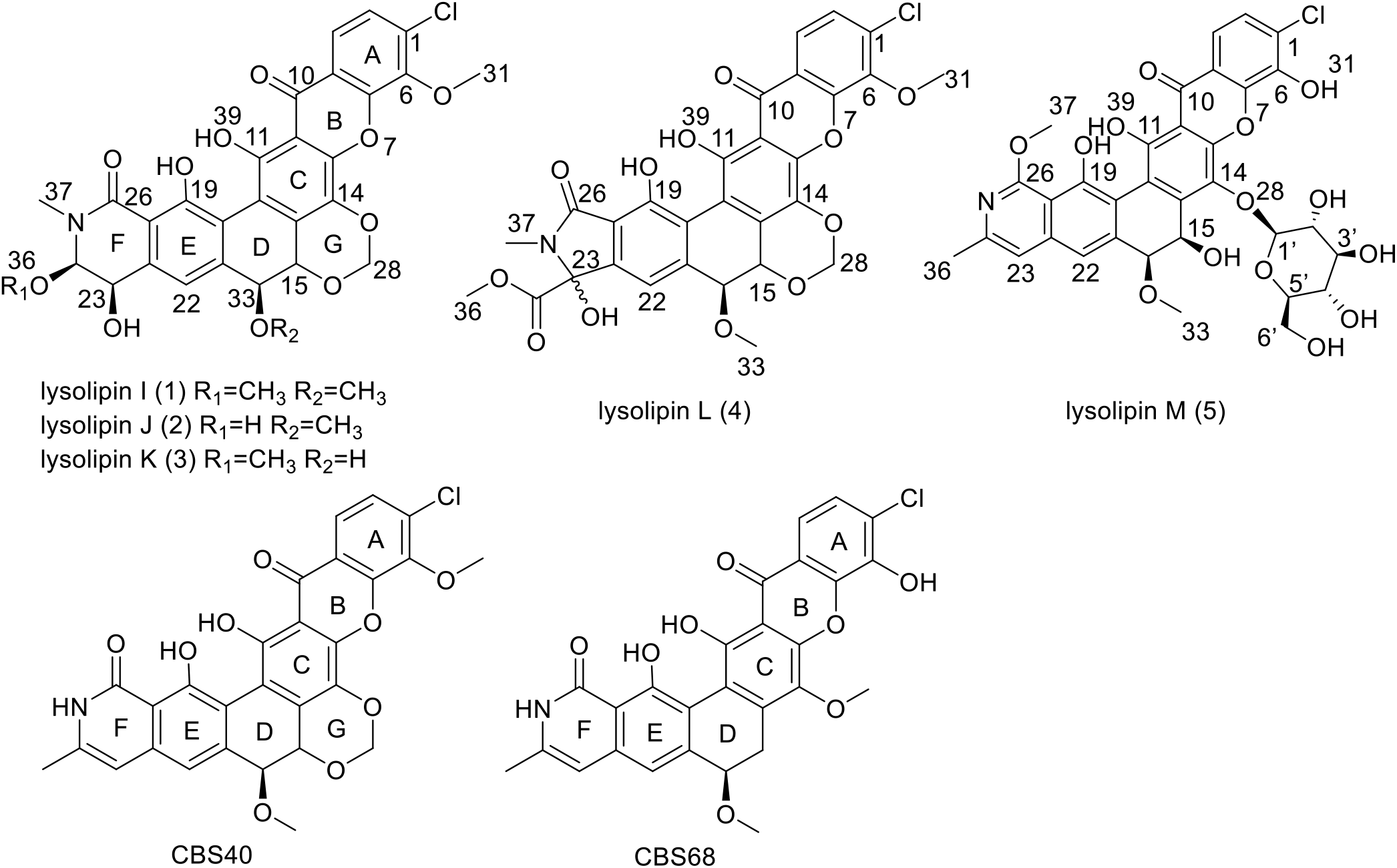
Structures of lysolipins I–M (**1**–**5**) isolated from *Streptomyces* sp. P8-2B18 and lysolipin analogues (CBS40 and CBS68) generated by genetic engineering^10^ or chemical modifications.^9^

Compound **1** was isolated as yellow needle-like crystals with a molecular formula of C_29_H_24_NO_11_Cl deduced by high-resolution ESIMS. The ^1^H NMR revealed signals for three *sp*^*2*^ aromatic protons (*δ*_H_ 7.97, 7.40, 7.10), four *sp*^*3*^ oxymethine protons (*δ*_H_ 5.04, 5.02, 4.72, 4.47), one methyl singlet (*δ*_H_ 3.36), and three methoxy protons (*δ*_H_ 4.17, 3.56, 3.35) along with one methylene *sp*^2^ upfield doublet protons at (*δ*_H_ 5.41, 5.64). Moreover, two downfield signals at *δ*_H_ 12.98 and *δ*_H_ 13.19 indicate the presence of two chelated protons. HSQC and HMBC spectra of **1** showed 29 carbon resonances that were attributed to one ketone carbonyl (*δ*_C_ 181.9), an amide/ester carbonyl (*δ*_C_ 168.5), three methoxy carbons (*δ*_C_ 61.9, 58.0, 57.6), a methyl group (*δ*_C_ 36.6), four oxymethines (*δ*_C_ 92.0, 78.9, 75.4, 67.8), a dioxymethylene (*δ*_C_ 91.1), three aromatic carbons (*δ*_C_ 125.4, 120.6, 115.9), and 15 quaternary carbons (**Table 1**).

**Table 1.**
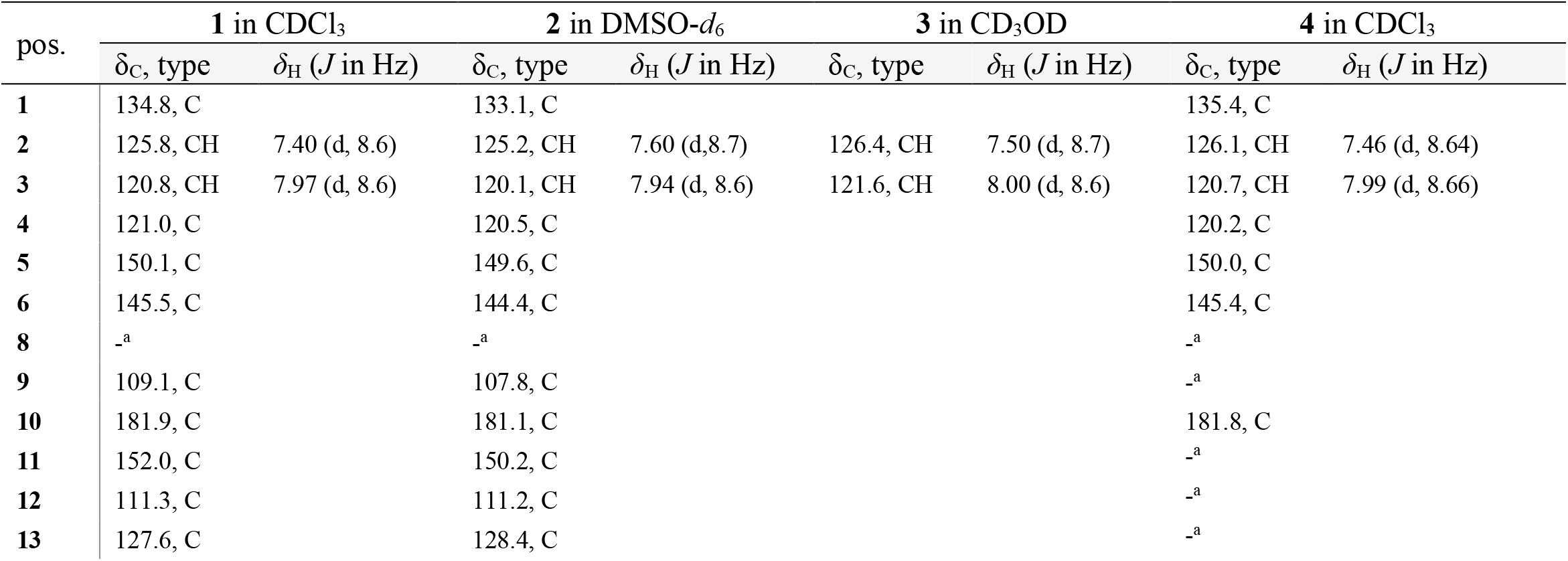
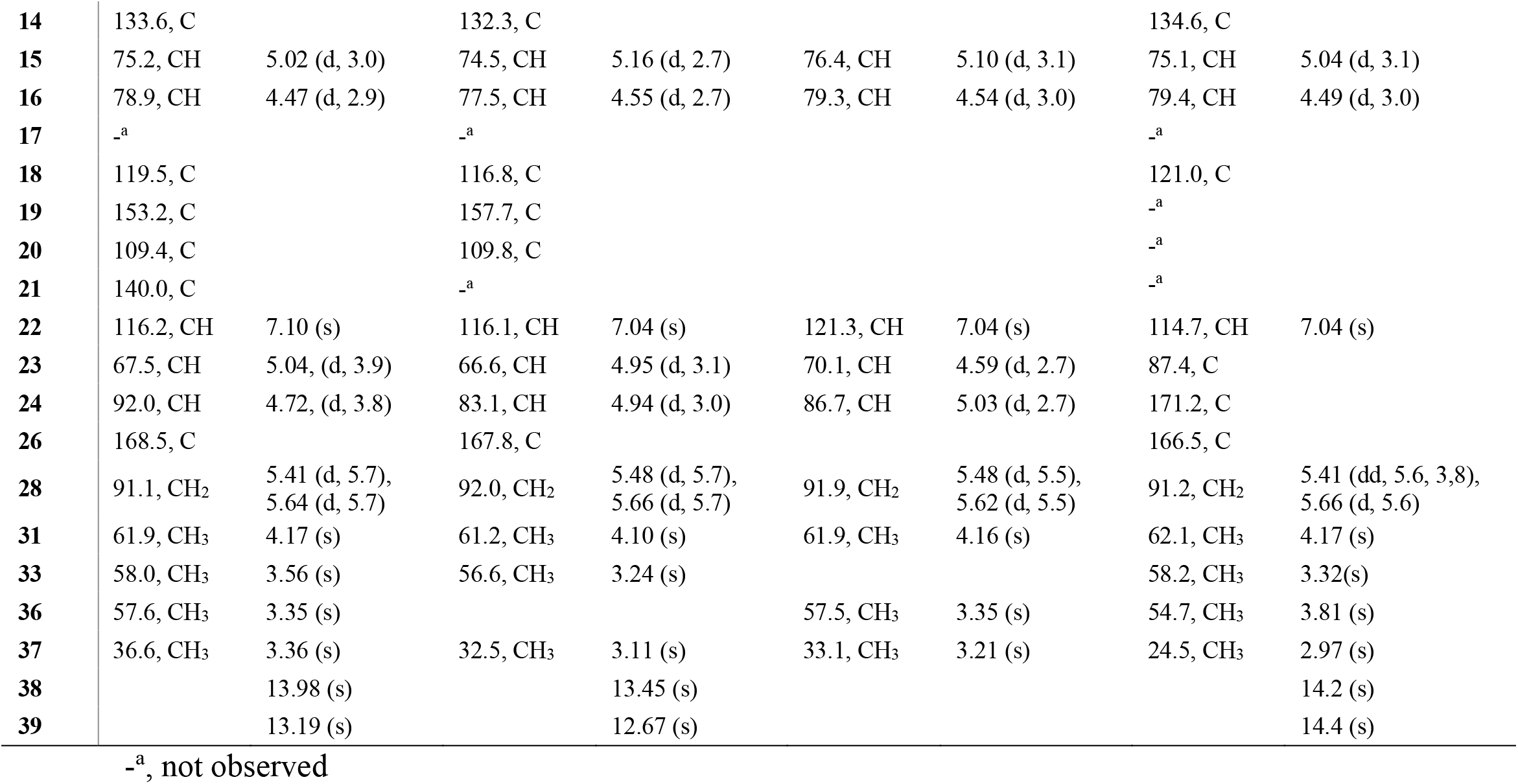
^1^H (800 MHz) and ^13^C (200 MHz) NMR Data for Lysolipins I–L (**1**–**4**).

| pos. | 1 in CDCl<sub>3</sub> |  | 2 in DMSO-*d*<sub>6</sub> |  | 3 in CD<sub>3</sub>OD |  | 4 in CDCl<sub>3</sub> |  |
| --- | --- | --- | --- | --- | --- | --- | --- | --- |
| | $\delta_{\text{C}}$, type | $\delta_{\text{H}}$ (*J* in Hz) | $\delta_{\text{C}}$, type | $\delta_{\text{H}}$ (*J* in Hz) | $\delta_{\text{C}}$, type | $\delta_{\text{H}}$ (*J* in Hz) | $\delta_{\text{C}}$, type | $\delta_{\text{H}}$ (*J* in Hz) |
| **1** | 134.8, C |  | 133.1, C |  |  |  | 135.4, C |  |
| **2** | 125.8, CH | 7.40 (d, 8.6) | 125.2, CH | 7.60 (d, 8.7) | 126.4, CH | 7.50 (d, 8.7) | 126.1, CH | 7.46 (d, 8.64) |
| **3** | 120.8, CH | 7.97 (d, 8.6) | 120.1, CH | 7.94 (d, 8.6) | 121.6, CH | 8.00 (d, 8.6) | 120.7, CH | 7.99 (d, 8.66) |
| **4** | 121.0, C |  | 120.5, C |  |  |  | 120.2, C |  |
| **5** | 150.1, C |  | 149.6, C |  |  |  | 150.0, C |  |
| **6** | 145.5, C |  | 144.4, C |  |  |  | 145.4, C |  |
| **8** | –<sup>a</sup> |  | –<sup>a</sup> |  |  |  | –<sup>a</sup> |  |
| **9** | 109.1, C |  | 107.8, C |  |  |  | –<sup>a</sup> |  |
| **10** | 181.9, C |  | 181.1, C |  |  |  | 181.8, C |  |
| **11** | 152.0, C |  | 150.2, C |  |  |  | –<sup>a</sup> |  |
| **12** | 111.3, C |  | 111.2, C |  |  |  | –<sup>a</sup> |  |
| **13** | 127.6, C |  | 128.4, C |  |  |  | –<sup>a</sup> |  |
| **14** | 133.6, C |  | 132.3, C |  |  |  | 134.6, C |  |
| **15** | 75.2, CH | 5.02 (d, 3.0) | 74.5, CH | 5.16 (d, 2.7) | 76.4, CH | 5.10 (d, 3.1) | 75.1, CH | 5.04 (d, 3.1) |
| **16** | 78.9, CH | 4.47 (d, 2.9) | 77.5, CH | 4.55 (d, 2.7) | 79.3, CH | 4.54 (d, 3.0) | 79.4, CH | 4.49 (d, 3.0) |
| **17** | -<sup>a</sup> |  | -<sup>a</sup> |  |  |  | -<sup>a</sup> |  |
| **18** | 119.5, C |  | 116.8, C |  |  |  | 121.0, C |  |
| **19** | 153.2, C |  | 157.7, C |  |  |  | -<sup>a</sup> |  |
| **20** | 109.4, C |  | 109.8, C |  |  |  | -<sup>a</sup> |  |
| **21** | 140.0, C |  | -<sup>a</sup> |  |  |  | -<sup>a</sup> |  |
| **22** | 116.2, CH | 7.10 (s) | 116.1, CH | 7.04 (s) | 121.3, CH | 7.04 (s) | 114.7, CH | 7.04 (s) |
| **23** | 67.5, CH | 5.04, (d, 3.9) | 66.6, CH | 4.95 (d, 3.1) | 70.1, CH | 4.59 (d, 2.7) | 87.4, C |  |
| **24** | 92.0, CH | 4.72, (d, 3.8) | 83.1, CH | 4.94 (d, 3.0) | 86.7, CH | 5.03 (d, 2.7) | 171.2, C |  |
| **26** | 168.5, C |  | 167.8, C |  |  |  | 166.5, C |  |
| **28** | 91.1, CH<sub>2</sub> | 5.41 (d, 5.7),<br>5.64 (d, 5.7) | 92.0, CH<sub>2</sub> | 5.48 (d, 5.7),<br>5.66 (d, 5.7) | 91.9, CH<sub>2</sub> | 5.48 (d, 5.5),<br>5.62 (d, 5.5) | 91.2, CH<sub>2</sub> | 5.41 (dd, 5.6, 3.8),<br>5.66 (d, 5.6) |
| **31** | 61.9, CH<sub>3</sub> | 4.17 (s) | 61.2, CH<sub>3</sub> | 4.10 (s) | 61.9, CH<sub>3</sub> | 4.16 (s) | 62.1, CH<sub>3</sub> | 4.17 (s) |
| **33** | 58.0, CH<sub>3</sub> | 3.56 (s) | 56.6, CH<sub>3</sub> | 3.24 (s) |  |  | 58.2, CH<sub>3</sub> | 3.32(s) |
| **36** | 57.6, CH<sub>3</sub> | 3.35 (s) |  |  | 57.5, CH<sub>3</sub> | 3.35 (s) | 54.7, CH<sub>3</sub> | 3.81 (s) |
| **37** | 36.6, CH<sub>3</sub> | 3.36 (s) | 32.5, CH<sub>3</sub> | 3.11 (s) | 33.1, CH<sub>3</sub> | 3.21 (s) | 24.5, CH<sub>3</sub> | 2.97 (s) |
| **38** |  | 13.98 (s) |  | 13.45 (s) |  |  |  | 14.2 (s) |
| **39** |  | 13.19 (s) |  | 12.67 (s) |  |  |  | 14.4 (s) |

-<sup>a</sup>, not observed

Analysis of COSY spectrum revealed three spin systems that included H-2/H-3, H-15/H-16, and H-23/H-24. The HMBC correlations (**Figure 2**) from *δ*_H_ 7.97 (H-3) to *δ*_C_ 181.9 (C-10) and 150.1 (C-5) confirmed the connection of 1,4,5,6-tetrasubstituted aromatic system (ring A) to ring B, while both H-2 and H-3 showed HMBC correlations to a relatively upfield carbon at *δ*_C_ 134.8 (C-1) bearing a chloride atom. The attachment of ring D to E was supported by the HMBC correlations from *δ*_H_ 4.47 (H-16) to the aromatic carbon *δ*_C_ 116.2 (C-22). The structure of ring F was established by the HMBC correlations of H-23 (*δ*_H_ 5.04) to the aromatic quaternary carbon C-21 (*δ*_C_ 140.0), the methyl protons (OCH_3_-36 and NCH_3_-37) to C-24, as well as NCH_3_-37 to C-26. Further HMBC spectrum analysis allowed the assignment of ring G based on the correlations of the acetal protons (H-28) to the oxymethine carbon *δ*_C_ 75.2 (C-15) and the quaternary carbon *δ*_C_ 133.6 (C-14). Compound **1** was then identified as lysolipin I as its NMR data are in good agreement with those reported in the literature.^3^ Confirmation of the absolute configuration of lysolipin I was further deduced from the positive specific rotation.

**Figure 2.**
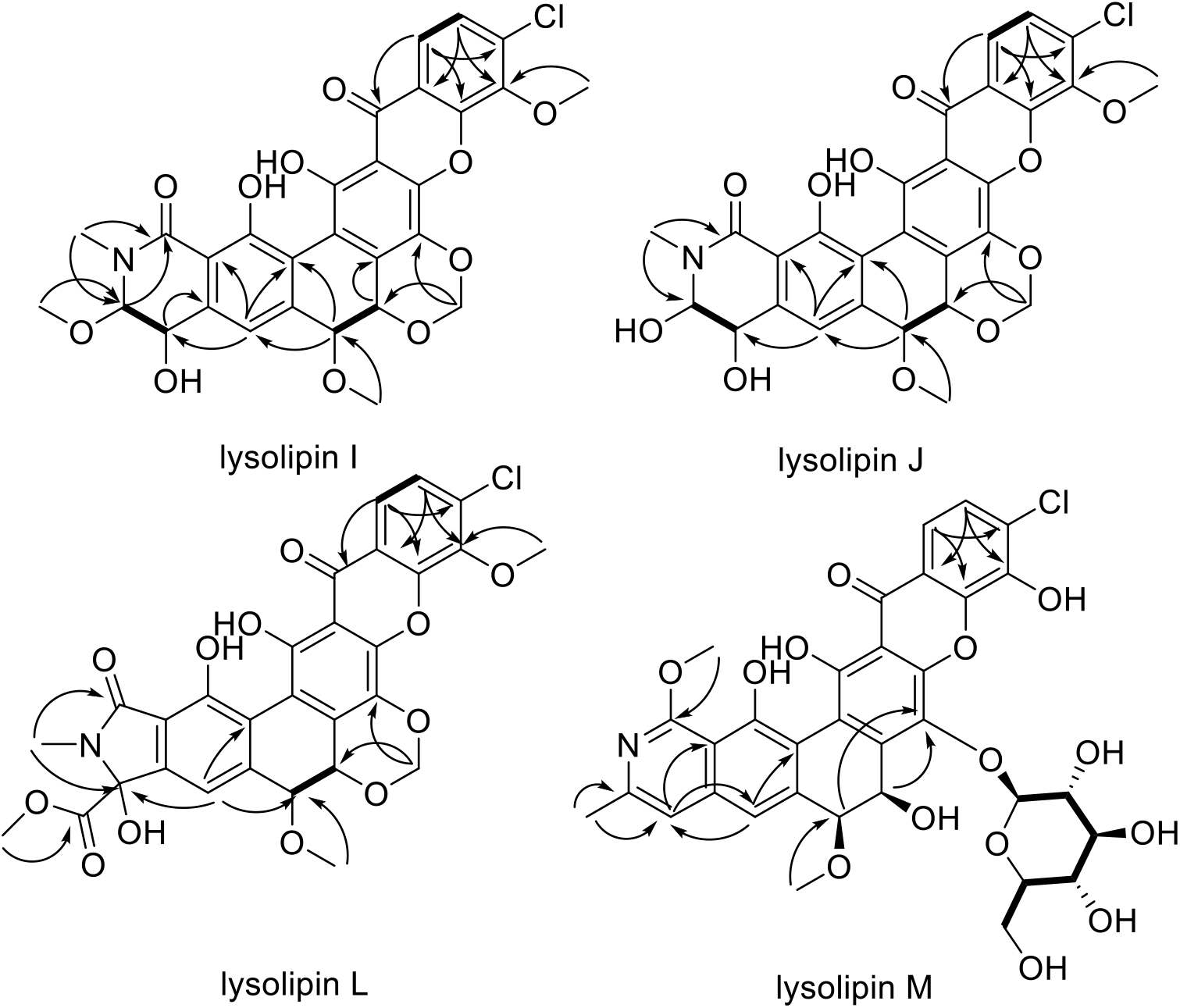
^1^H-^1^H COSY (**⁃**) and HMBC (↷) correlations of lysolipins I, J, L, and M (**1, 2, 4**, and **5**).

Compounds **2** and **3** were isolated as derivatives of lysolipin I, with only difference of one less methyl group compared to **1**. The molecular formula of C_28_H_22_NO_11_Cl was deduced by HRESIMS data.

In compound **2**, the ^1^H NMR data revealed similar signals for three *sp*^*2*^ aromatic protons (*δ*_H_ 7.94, 7.60, 7.13), four *sp*^*3*^ oxymethine protons (*δ*_H_ 5.16, 4.95, 4.94, 4.55), one methyl singlet (*δ*_H_ 3.21), and only two methoxy groups protons (*δ*_H_ 3.11, 3.24) along with two doublet protons of the methylene dioxybridge (*δ*_H_ 5.48, 5.66). The COSY and HMBC correlations (**Figure 2**) revealed a similar core structure to lysolipin I. The only difference is the absence of an oxygenated methyl group which was connected to C-24 in lysolipin I. Thus, compound **2** herein was named lysolipin J.

Similar to lysolipin J, the NMR data of compound **3** revealed a similar core structure to lysolipin I. The only difference is the absence of an oxygenated methyl group which was connected to C-16 in lysolipin I, while the methoxy group connected to C-24 (*δ*_H_ 3.35, *δ*_C_ 57.5) was conserved. This led to the confirmation of the structure of **3** as lysolipin K.

Compound **4** was isolated as another variant of lysolipin I, with a molecular formula of C_29_H_22_ClNO_12_. This indicated one more DBE compared to lysolipin I. In the ^1^H NMR spectrum, compound **4** exhibited similar signals as lysolipin I except the absence of signals for H-23 and H-24. This indicated rearrangement in ring F compared to lysolipin I. Signals from ^13^C NMR spectrum showed a downfield oxygenated quaternary carbon (*δ*_C_ 87.4, C-23) and an extra carbonyl group (*δ*_C_ 171.2, C-24). The position of C-23 and C-24 was confirmed through the HMBC correlations of H-22 (*δ*_H_ 7.96) with C-23, OMe-36 (*δ*_H_ 3.81) with C-24 (**Figure 2**). Moreover, the methyl group *δ*_H_ 2.97 (C-37) showed correlations with C-26 (*δ*_C_ 166.5) and C-23. Those key HMBC correlations led to a proposed five-membered lactam ring F. The resulting structure featuring a hexacyclic xanthone core and 5-membered heterocyclic ring is unprecedented for reported polycyclic xanthones. Interestingly, the NMR spectra revealed two similar groups of signals, which indicated compound **4** existed as a mixture of two diastereomers. From biosynthesis perspective, compound **4** was proposed to derive from compound **2** through a spontaneous Favorskii Rearrangement^15^ (**Figure 3**), without stereochemical selectivity during the rearrangement and formation of the chiral center (C-23). Accordingly, compound **4** was determined to be lysolipin L. Instead of a six-membered lactam ring in lysolipins I, J and K, lysolipin L featured a five-membered lactam F ring.

**Figure 3.**
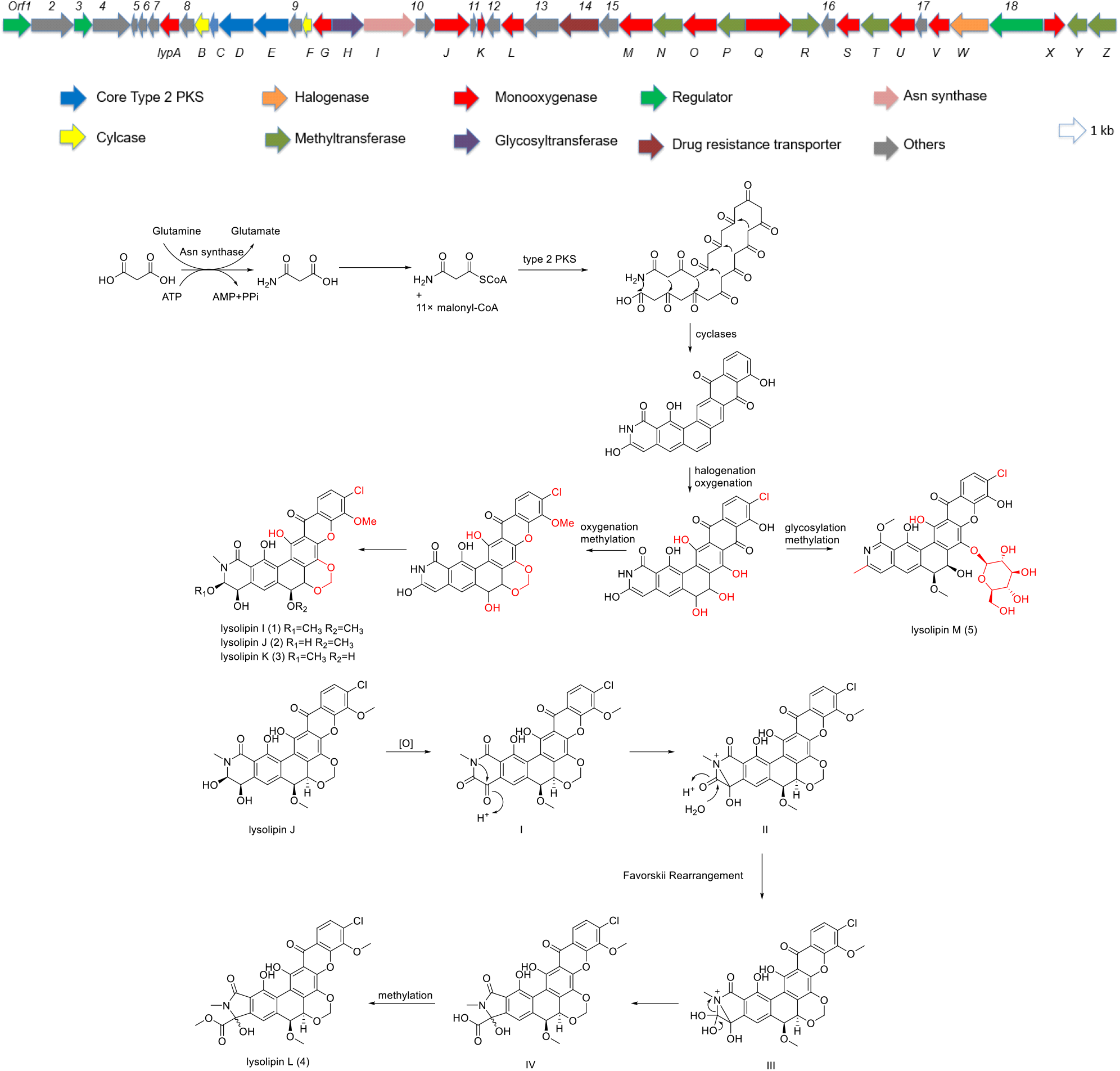
Proposed biosynthetic pathway of lysolipins. Proposed annotation of individual ORFs can be found in **Table S6**, supplemental information.

Compound **5** was isolated as a glycosylated lysolipin variant. It appeared to be more hydrophilic as it was eluted at a relatively early retention time compared to the other lysolipins. It has a molecular formula of C_33_H_30_NO_14_Cl, suggested the presence of an extra sugar moiety, which was confirmed by the MS/MS fragmentation (**Figure S27**). The ^1^H NMR spectrum showed the presence of four aromatic proton signals including two doublets *δ*_H_ 7.61(H-2), *δ*_H_ 7.97 (H-3) and two singlets *δ*_H_ 7.15 (H-22), *δ*_H_ 6.51 (H-23) (**Table 2**). Similar ring A moiety was confirmed by COSY correlations between H-2 and H-3, as well as HMBC correlations from H-2 to quaternary carbon C-4 (*δ*_C_ 120.1) and phenyl carbon C-6 (*δ*_C_ 140.6), and from H-3 to C-1 (*δ*_C_ 133.8) and C-5 (*δ*_C_ 145.0). The same structure of ring D was confirmed by COSY correlation between H-15 (*δ*_H_ 5.36) and H-16 (*δ*_H_ 4.25), together with the HMBC correlations from H-15 and H-16 to C-14 (*δ*_C_ 143.2). To confirm the remaining ring structures (ring E and F), key HMBC correlations (**Figure 2**) are between H-22 (*δ*_H_ 7.15) with C-23 (*δ*_C_ 110.6), and H-23 (*δ*_H_ 6.51) with C-22 (*δ*_C_ 105.1). The methoxy proton H-37 (*δ*_H_ 4.08) showed HMBC correlation to C-26 (*δ*_C_ 136.6). Furthermore, HMBC correlations from the methyl protons H-36 (*δ*_H_ 2.26) to C-23 and C-24 supported ring F as an aromatic pyridine ring, which has been reported in CBS40 and CBS68 (**Figure 1**).

**Table 2.**
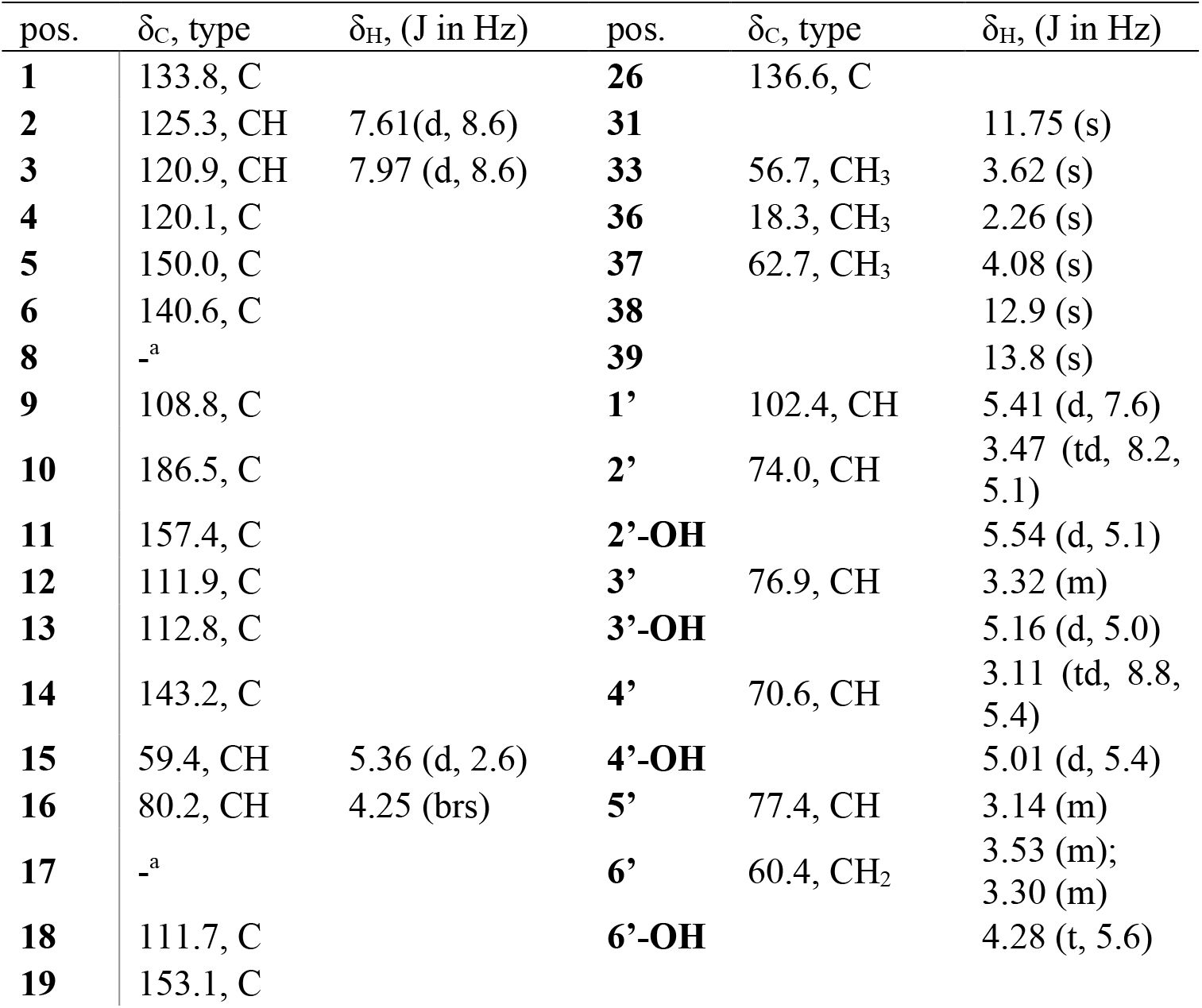
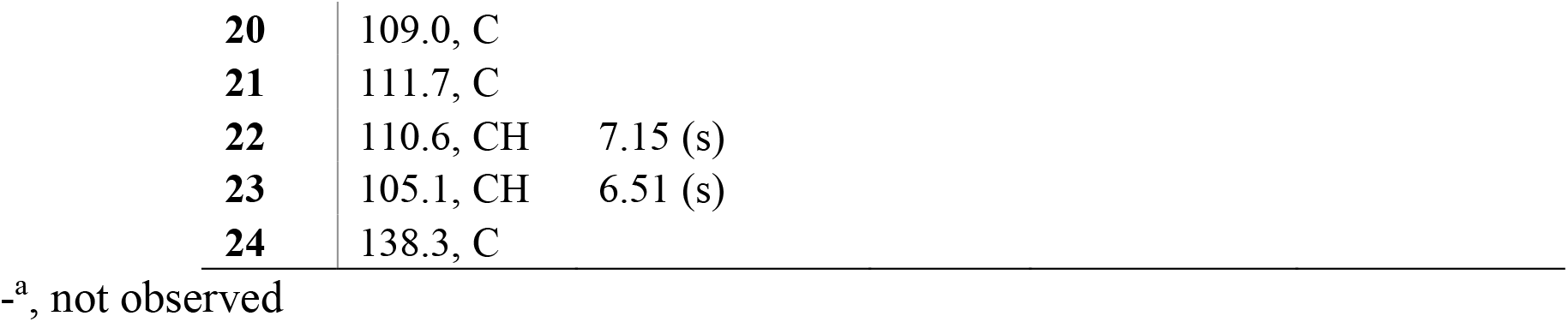
^1^H (800 MHz) and ^13^C (200 MHz) NMR Data for Lysolipin M (**5**) in DMSO-*d*_*6*_.

| pos. | δ<sub>C</sub>, type | δ<sub>H</sub>, (J in Hz) | pos. | δ<sub>C</sub>, type | δ<sub>H</sub>, (J in Hz) |
| --- | --- | --- | --- | --- | --- |
| **1** | 133.8, C |  | **26** | 136.6, C |  |
| **2** | 125.3, CH | 7.61(d, 8.6) | **31** |  | 11.75 (s) |
| **3** | 120.9, CH | 7.97 (d, 8.6) | **33** | 56.7, CH<sub>3</sub> | 3.62 (s) |
| **4** | 120.1, C |  | **36** | 18.3, CH<sub>3</sub> | 2.26 (s) |
| **5** | 150.0, C |  | **37** | 62.7, CH<sub>3</sub> | 4.08 (s) |
| **6** | 140.6, C |  | **38** |  | 12.9 (s) |
| **8** | -<sup>a</sup> |  | **39** |  | 13.8 (s) |
| **9** | 108.8, C |  | **1'** | 102.4, CH | 5.41 (d, 7.6) |
| **10** | 186.5, C |  | **2'** | 74.0, CH | 3.47 (td, 8.2,<br>5.1) |
| **11** | 157.4, C |  | **2'-OH** |  | 5.54 (d, 5.1) |
| **12** | 111.9, C |  | **3'** | 76.9, CH | 3.32 (m) |
| **13** | 112.8, C |  | **3'-OH** |  | 5.16 (d, 5.0) |
| **14** | 143.2, C |  | **4'** | 70.6, CH | 3.11 (td, 8.8,<br>5.4) |
| **15** | 59.4, CH | 5.36 (d, 2.6) | **4'-OH** |  | 5.01 (d, 5.4) |
| **16** | 80.2, CH | 4.25 (brs) | **5'** | 77.4, CH | 3.14 (m) |
| **17** | -<sup>a</sup> |  | **6'** | 60.4, CH<sub>2</sub> | 3.53 (m);<br>3.30 (m) |
| **18** | 111.7, C |  | **6'-OH** |  | 4.28 (t, 5.6) |
| **19** | 153.1, C |  |  |  |  |

|  |  |  |
| --- | --- | --- |
| 20 | 109.0, C |  |
| 21 | 111.7, C |  |
| 22 | 110.6, CH | 7.15 (s) |
| 23 | 105.1, CH | 6.51 (s) |
| 24 | 138.3, C |  |

-<sup>a</sup>, not observed

The remaining ^1^H NMR and ^13^C NMR signals have been established as a hexose moiety through COSY and HMBC correlations (**Figure 2**). COSY correlations of H-1’ (*δ*_H_ 5.41)/H-2’ (*δ*_H_ 3.47)/H-3’ (*δ*_H_ 3.32)/H-4’ (*δ*_H_ 3.11)/H-5’ (*δ*H 3.14)/H_2_-6’ (*δ*_H_ 3.30 and 3.53) and the large ^3^*J*_H1’-H2’_ value with a relatively high carbon shift (*δ*_C_ 102.4) shows a *β* configuration at the anomeric position (CH-1’). The large coupling constants observed between H-1’ and H-2’ (7.6 Hz), H-2’ and H-3’ (8.2 Hz), H-3’ and H-4’ (8.8 Hz), and H-4’ and H-5’ (8.8 Hz) placed the hydroxy groups at C-2’, C-3’ and C-4’, as well as the CH_2_OH-6’ group in an equatorial position, making substructure a *β*-glucose^16^. However, the absolute configuration of the sugar could not be determined because of insufficient material for hydrolysis and comparison with authentic standards. Considering the relative downfield chemical shift (*δ*_C_ 143.2) for C-14 (*δ*_C_ 133.6 in **1**) and the absence of C-28 indicated the disruption of ring G, we suggested the attachment of a sugar moiety to the at C-14. Chemical shifts for the other three phenyl groups (C-6, C-11 and C-19) remained conserved compared to compounds **1–4**. Compound **5** was confirmed as a glycosylated analogue of lysolipin, named as lysolipin M.

Hybrid whole-genome sequencing and assembly of the *Streptomyces* sp. P8-2B18 resulted in a complete genome of one chromosome and two plasmids totalling 8,773,358 bp. The estimated Average Nucleotide Identity (ANI) according to autoMLST2^17^ is 94.8% and the digital DNA-DNA hybridization (DDH) calculated via TYGS^18^ is 52%, with the closely related species *Streptomyces kronopolitis*. Following antiSMASH analysis,^19^ the genome contains 41 predicted BGCs, including a 72,540 bp T2 PKS matching the lysolipin I BGC (**Table S5**), at 5,273,738–5,346,277 on the chromosome. This cluster is highly similar to the previously described lysolipin BGCs of *Streptomyces tendae* Tue 4042^20^ (**Figure 4**). However, comparative genomic analysis on *Streptomyces* sp. P8-2B18 with the known lysolipin producer *Streptomyces tendae* revealed that they are phylogenetically distinct (**Figure S30**).

**Figure 4.**
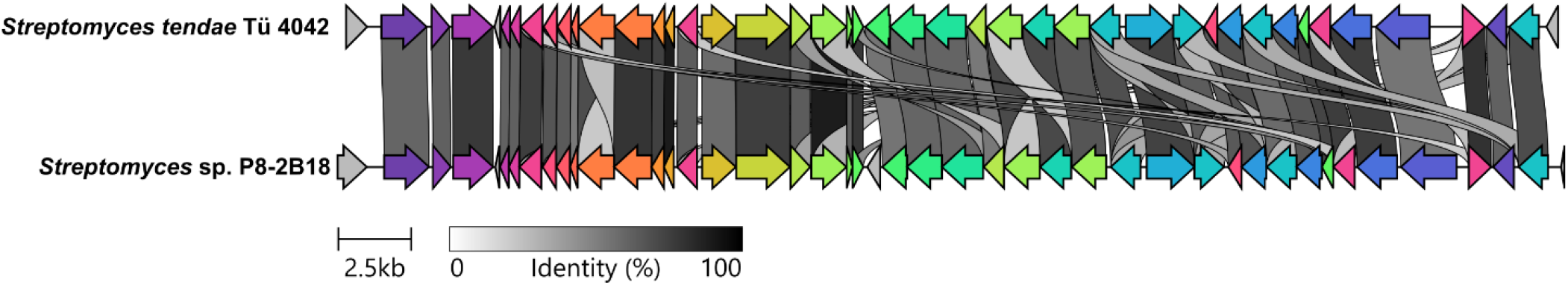
Clinker^21^ comparisons of the lysolipin biosynthetic gene clusters in *Streptomyces tendae* Tue 4042 (producer of lysolipin I) and *Streptomyces* sp. P8-2B18 (producer of lysolipins I–M). The alignment with gene coloring based on predicted or experimentally elucidated involvement in biosynthesis.

Biosynthesis of lysolipin has been proposed in *Streptomyces tendae*.^22^ The core type 2 PKS, cyclase, halogenase, oxygenases, methyltransferase as well as the glycosyltransferase (LypH) are conserved in both producers. For the first time, we obtained lysolipin M as a glycosylated version of lysolipin. Meanwhile, one of the oxygenases facilitated the cascade of Favorskii rearrangement, leading to the formation of lysolipin L with a five-membered lactam F ring. Interestingly, lysolipin M has an extra methyl group (Me-36) at C-24 position. This is likely formed by one of the six methyl transferases (LypN, P, R, T, Y and Z). Here discovery of lysolipins I–M highlighted versatility of the *lyp* BGC and chemical diversity of the lysolipin family. The proposed biosynthetic pathway of lysolipins I–M can be seen in **Figure 3**.

In summary, we have reported four new variants of lysolipins, lysolipins J-M. While lysolipin J and K are close derivatives of lysolipin I, lysolipin L and M represent novel types of lysolipins. Lysolipin L features a five-membered lactam F ring, which was unprecedented in reported lysolipins. Lysolipin M has a novel skeleton, with an extra methyl (Me-36) and a glycosyl group compared to a 1,3-oxane ring in lysolipin I. To gain insight into their biosynthesis, we have looked into the *lyp* BGC. Based on the shared biosynthetic pathway, we proposed the same absolute configurations at C-15 and C-16 for all lysolipins. Similar ECD curves were observed for lysolipin I, L and M (Figure S28, supplemental information).

Many polycyclic xanthones have been reported as potent cytotoxic agents. For example, xantholipins A and B exhibited cytotoxicity against five human cancer cell lines with IC_50_ values in the sub-micromolar to nanomolar range.^2^ We tested the effects of lysolipins I, J, K, L, and M on cell viability and proliferation in the human epithelial prostate cancer cell lines LNCaP and C4-2B. Compounds showing antiproliferative effect were further evaluated in multi-drug resistant sublines, LNCaP^R^ and C4-2B^R^, which are characterized by overexpression of the drug efflux pump P-glycoprotein (Pgp). To assess the antiproliferative activity, we monitored cell confluency over time in parental prostate cancer cell lines (LNCaP and C4-2B) and their multidrug-resistant (MDR) derivatives (LNCaP^R^ and C4-2B^R^) following treatment with 10 μM of each compound. Vehicle-treated controls exhibited robust proliferation, reaching >90% confluency by the end of the assay. In contrast, lysolipins I, J, and K produced near-complete growth inhibition in LNCaP and C4-2B cells, maintaining confluency levels below 5% throughout the experiment. Lysolipins L and M showed weak to moderate activity, with confluency increasing to 35–60%. Docetaxel (100 nM) was used as a cytotoxic control.

In the MDR sublines, lysolipins remained active, albeit with slightly reduced potency. Lysolipin I was the most effective, limiting confluency to ∼10% in C4-2BR and <5% in LNCaPR. Lysolipins J and K allowed partial growth, with cell confluency reaching ∼20% in C4-2BR and 8–10% in LNCaPR. Lysolipin M exhibited weak inhibition in resistant cells (50–60% confluency). These findings indicate that lysolipins I–K are highly potent inhibitors of prostate cancer cell proliferation, although their efficacy is diminished in MDR cells, consistent with P-glycoprotein-mediated drug efflux.

To further assess potency, IC_50_ values were estimated using a sigmoidal model based on cell viability after 48h exposure of LNCaP and C4-2B parental cells to a concentration series (0, 0.01, 0.1, 0.2, 1, and 10µM) of lysolipins I-M. Lysolipins I–K exhibited submicromolar IC_50_ values in parental lines, with lysolipin I being the most potent (0.20 μM in LNCaP and C4-2B), followed by lysolipin J (0.31 μM and 0.28 μM in LNCaP and C4-2B, respectively) and lysolipin K (0.53 μM and 0.61 μM in LNCaP and C4-2B, respectively). In contrast, lysolipins L and M showed markedly lower potency (>10 μM and 5.6 μM, respectively). These data confirm that lysolipins I–K are highly potent cytotoxic agents and may represent interesting preliminary leads. The reduced activity observed in multidrug-resistant sublines suggests potential susceptibility to P-glycoprotein-mediated efflux, which could compromise efficacy in resistant tumors. Moreover, the selectivity of these compounds for cancer versus non-malignant cells remains to be established, and insufficient selectivity could limit their therapeutic window due to potential toxicity in normal tissues. Structural optimization aimed at improving selectivity and reducing efflux liability may be required before these compounds can be viable anticancer drug candidates.

The significant cytotoxicity demonstrated by polycyclic xanthones documented in literature and verified in our recent research, indicates their potential as highly effective chemotherapeutic agents against cancer. However, despite this promising cytotoxicity, there remains a gap in our comprehension of the cytotoxic mechanism of Bioactive Polycyclic Xanthone Nanoparticles (BPXNPs). To address this knowledge gap, we conducted computational studies, particularly molecular docking, to uncover potential molecular interactions and binding modes of BPXNPs with critical biomolecular targets associated with cancer. Herein, we propose that their inhibition of DNA synthesis leading to apoptosis by targeting topoisomerase IIβ. This enzyme regulates DNA topology by creating transient breaks in the DNA molecule. Etoposide, a chemotherapeutic poison targeting type II topoisomerases, stabilizes cleaved-DNA intermediates, resulting in permanent DNA breaks.^23^ Consequently, antiproliferative agents act as a bridge between the DNA helix and the protein, impeding their separation and inducing apoptosis. Additionally, during docking experiments of lysolipin with Topoisomerase IIβ (PDB 3QX3),^24^ the chelation of the cytotoxic compound between the two DNA strands was observed. Additionally, docking experiments of lysolipin with the Topoisomerase IIβ–DNA complex (PDB ID: 3QX3) revealed a stable binding mode in which the planar aromatic core of lysolipin is positioned between the two DNA strands at the enzyme–DNA cleavage site (**Figure 5**). This intercalative binding is further stabilized by π–π stacking interactions with the DNA base pairs, while the peripheral functional groups of lysolipin form hydrogen-bonding and electrostatic interactions with key amino acid residues lining the Topoisomerase IIβ active site. Notably, the predicted binding pose suggests that lysolipin bridges both DNA and protein components of the ternary complex, thereby potentially stabilizing the cleavage complex and preventing DNA religation. Such a binding mode is consistent with a Topoisomerase II poison–like mechanism and provides a structural rationale for the observed cytotoxic activity of lysolipin.^24,25^

**Figure 5.**
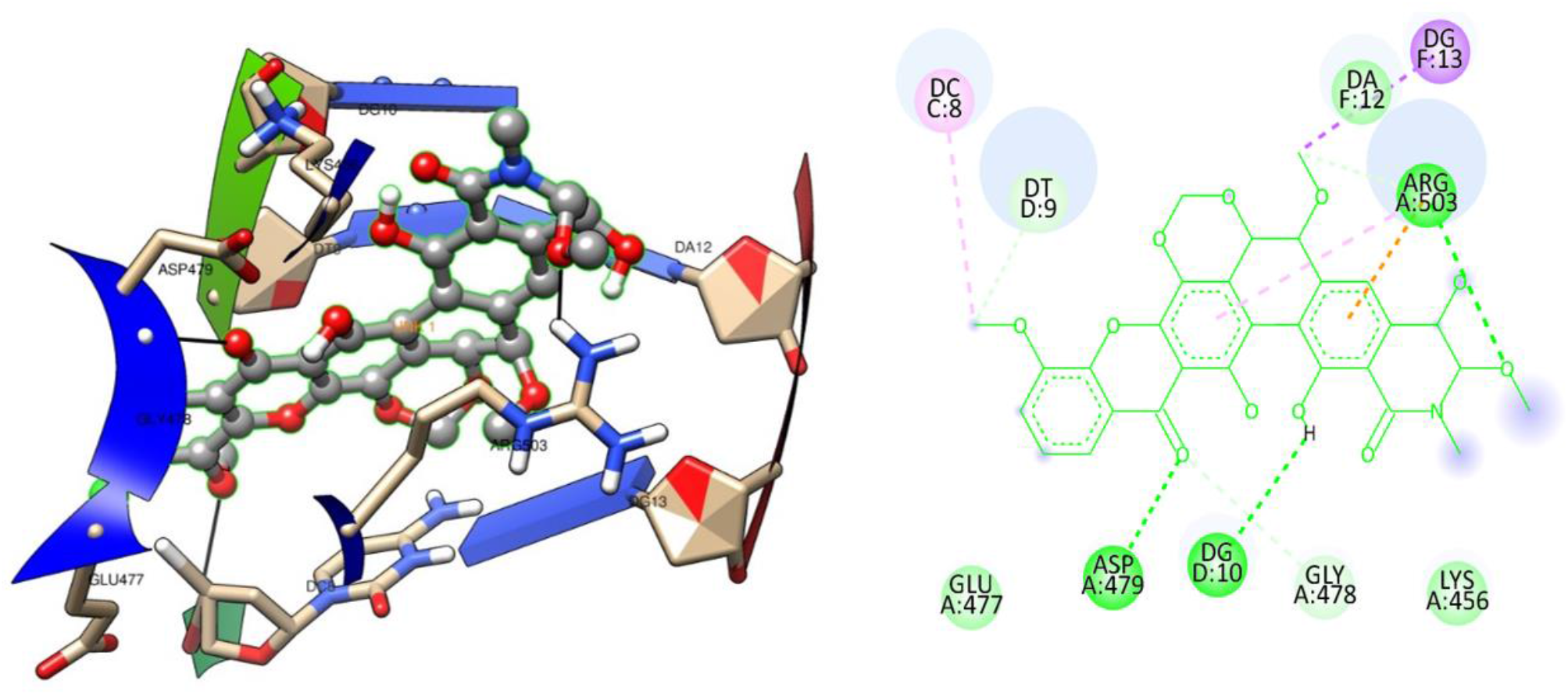
The binding orientation of lysolipin I (grey stick) inside the 3QX3 pocket.

While lysolipin I represents the most bioactive analogue, lysolipins J–K are less potent, but still in the same order of magnitude. Meanwhile, lysolipin M was much less active and lysolipin L lost activity. Based on the docking experiments of lysolipin I, ring F seems to be essential in the interactions and this explains why the other derivatives are less active. This is also reflected in the other lysolipin I derivatives (e.g., CBS40 and CBS68, **Figure 1**) generated by chemical or genetic engineering, where the cytotoxicity was dramatically reduced.^10^

Lysolipin I was reported to exhibit potent antibacterial activity in the low-nanomolar range against both Gram-positive and Gram-negative bacteria including multidrug-resistant strains.^3^ However, their cytotoxicity has hindered its applications as a new antibiotic. Efforts have been taken to generate analogues with less cytotoxicity.^9^ Nonetheless, no studies have been done to evaluate their activities against fungal pathogens. In this study, lysolipins (**1–5**) were evaluated against *Staphylococcus aureus* 8325 and *Aspergillus flavus* IBT30114. Lysolipin I displayed most potent activity with MIC values of 0.25 and 1 μg/mL against *S. aureus* and *A. flavus*, respectively, while lysolipins J and K with moderate activities. Modification in the ring F led to reduced activities in lysolipin L, whereas lysolipin M completely abolished activities ((**Table 3**). Despite their antimicrobial activities, the application as new antibiotics is challenged by their cytotoxicity. All the five isolated lysolipins donot exhibit selective activities in cytotoxicity or antimicrobial. Glycosylation led to a much lower cytotoxicity in lysolipin M. However, the antimicrobial activities were abolished at the same time. Although lysolipin L doesnot have cytotoxicity, it only exhibited moderate antimicrobial activities.

**Table 3.** Antimicrobial and cytotoxicity activities of isolated compounds.

| Test compound | MIC ($\mu\text{g/ml}$) values | | IC<sub>50</sub> values ($\mu\text{M}$) | |
| --- | --- | --- | --- | --- |
|  | *S. aureus* | *A. flavus* | LNCaP | C4-2B |
| lysolipin I (1) | 0.25 | 1 | 0.20 | 0.20 |
| lysolipin J (2) | 1 | 4 | 0.31 | 0.28 |
| lysolipin K (3) | 1 | 4 | 0.53 | 0.61 |
| lysolipin L (4) | 4 | 16 | >10 | >10 |
| lysolipin M (5) | $\geq 64$ | $\geq 64$ | 5.6 | 7.3 |
| tetracycline | 1 | - | - | - |
| amphotericin B | - | 4 | - | - |
| docetaxel | - | - | 0.00147 | 0.00206 |

## EXPERIMENTAL SECTION

### General Experimental Procedures

Optical rotations were recorded on an AUTOPOL III - S2 Dual Wavelength (589/546nm) Automatic Polarimeter ([Rudolph Research Analytical](https://www.labcompare.com/152-Rudolph-Research-Analytical/)). ECD spectra were obtained in MeOH using a 10 mm path cuvette on a JASCO J-1500 CD spectrometer. The NMR spectra were recorded on a Bruker AVANCE III 800 MHz spectrometer equipped with a 5 mm TCI CryoProbe using standard pulse sequences. The ^1^H and ^13^C NMR chemical shifts were reported with reference to the residual solvent signals at *δ*_H_ 7.26, *δ*_C_ 77.16 ppm for CDCl_3_, *δ*_H_ 3.35, *δ*_C_ 49.3 ppm for CD_3_OD, and *δ*_H_ 2.50, *δ*_C_ 39.52 for DMSO-*d*_6_. UHPLC-HRMS was performed on an Agilent Infinity 1290 UHPLC system equipped with a diode array detector. UV−vis spectra were recorded from 190 to 640 nm. All solvents and chemicals used for UHPLC-HRMS were LC-MS grade, while the solvents for extraction and chromatography were of HPLC grade. Water was purified using a Milli-Q system.

### Strain Isolation and Genome Sequencing

The bacterial strain was isolated from soil collected in 2020 from the UNESCO World Heritage Site, Jægersborg Deer Park (Dyrehaven in Danish), Denmark. The *Streptomyces* sp. P8-2B18 strain was inoculated in 50 mL of sterile ISP2 liquid medium (yeast extract 4.0 g, malt extract 10.0 g, and dextrose 4.0 g in 1.0 L of distilled water, Ph = 7.2) in a 300 mL flask, incubated at 28 °C and 160 rpm for 3 days. The genomic DNA was isolated and purified using a QIAGEN Genomic-tip G100 kit.^26^ The genomic DNA sequencing was performed using both Oxford Nanopore GridION and Illumina NovaSeq X Plus Series. First the nanopore reads were assembled with Flye v2.9.3 and then polished with Illumina reads using polypolish v0.6.1, all with standard settings. Taxonomy was inferred directly from nanopore reads with sylph, along with assembly-level classification with TYGS^18^ and autoMLST2^17^, the ladder of which was also used for phylogenetic inference. Biosynthetic gene clusters were found by submission to the antiSMASH online server^19^, and clinker^21^ was used to compare the lysolipin BGC with *S. tendae*.

### Data-Dependent LC-ESI-HRMS/MS Analysis

Ultra-high-performance liquid chromatography−diode array detection−quadrupole time-of-flight mass spectrometry (UHPLC-DAD-QTOFMS) was performed on an Agilent Infinity 1290 UHPLC system equipped with a diode array detector. Separation was achieved on a 250 × 2.1 mm i.d., 2.7 μm, Poroshell 120 phenyl-hexyl column (Agilent Technologies) held at 40 °C. The sample, was eluted at a flow rate of 0.35 mL min^−1^ using a linear gradient from 40% to 50% MeCN in Milli-Q water buffered with 20 mM formic acid over 20 min by an Agilent Infinity 1290 HPLC-DAD (Agilent Technologies) system. Mass spectrometry (MS) detection was performed on an Agilent 6545 QTOF MS equipped with an Agilent dual jet stream electrospray ion source (ESI) with a drying gas temperature of 160 °C, a gas flow of 13 L min^−1^, a sheath gas temperature of 300 °C, and a flow rate of 16 L min^−1^. Capillary voltage was set to 4000 V and nozzle voltage to 500 V in positive mode. MS spectra were recorded as centroid data, at an m/z of 100−1700, and auto MS/HRMS fragmentation was performed at three collision energies (10, 20, and 40 eV), on the three most intense precursor peaks per cycle. The acquisition rate was 10 spectra s^−1^. Data were handled using Agilent MassHunter Qualitative Analysis software (Agilent Technologies). Lock mass solution in 70% MeOH in water was infused in the second sprayer using an extra LC pump at a flow rate of 15 μL/min using a 1:100 splitter. The solution contained 1 μM tributylamine (SigmaAldrich) and 10 μM hexakis (2,2,3,3-tetrafluoropropoxy) phosphazene (Apollo Scientific Ltd., Cheshire, UK) as lock masses. **GNPS Workflow**. The acquired MS/MS data from analysis by liquid chromatography−diode array detection−quadrupole time-of flight mass spectrometry (LC-DAD-TOFMS) were preprocessed by MZmine2.53^27^ and analyzed using the feature-based molecular networking workflow on the Global Natural Product Social Molecular Networking platform^28^. The generated molecular networks were visualized in Cytoscape^29^.

### Fermentation, Extraction and Isolation

Liquid organic medium 79 (dextrose 10 g, bactopeptone10 g, casamino acids 1 g, yeast extract 2 g, NaCl 6 g, H_2_O 1 L; 2 × 100 mL/flask) was inoculated with a suspension of mycelium and spores (about 1 × 1 cm) of *Streptomyces* sp. P8-2B18 grown on agar slants or agar plates (ISP medium 2). After incubation for 48 h on a rotary shaker at 28 °C, the culture was transferred to 3200 mL of ISP2 (Yeast extract (Difco) 4.0 g, Malt extract (Difco) 10.0 g, Dextrose (Difco) 4.0 g, distilled water 1000.0 mL pH 7.2; in eight 1000 mL-scale Erlenmeyer flasks with 400 mL of medium ISP2 each) and incubated at 28 °C under shaking conditions for 48 h to yield the seed culture. The culture was then poured into a 75 L-scale fermenter filled with 50 L of medium ISP2 and fermented for 6 days. After fermentation, mycelia were lyophilized and sequentially extracted with methanol. The filtrated supernatant (40 L) was separated by an Amberchrom 161CGM resin LC column (10 × 10 cm, 1 L), and a linear gradient of H_2_O–MeOH (from 30% to 100% v/v, flow rate 0.18 L min–L, in 58 min) was applied to afford nine fractions (F1–F9).

The lyophilized fraction F7 (5 g) was loaded on normal-phase silica gel SNAP 100 g Biotage Flash Cartridge Isolera, the gradient used was DCM for 10 minutes and DCM/10%MeOH over 30 min (25 mL/min), finally DCM/50%MeOH for 10 min. The 10% MeOH subfraction (1g) was further fractionated using a linear gradient from 10–100 % MeOH on column RP-C_18_ HPLC to get seven subfractions (A–G). Lysolipin I (**1**; 10 mg), J (**2**; 2 mg), K (**3**; 2 mg) and L (**4**; 0.5 mg) were further purified from 7D on a phenyl-hexyl column (Luna, 100 Å, 250 × 10 mm, i.d., 5 μm, Phenomenex) held at 40 °C by linear gradient elution 50–80% to MeCN in Milli-Q water over 20 min.

Fractionation of subfraction 5 (3 g) was performed by SNAP 100 g column normal-phase silica gel (Grace, 15 μm/100 Å) flash column (Biotage, Uppsala, Sweden) using the Isolera One automated flash system (Biotage). The gradient used was DCM/10%MeOH over (45 mL/min). Fractions were automatically collected based on UV signals (254 and 320 nm). Followed by separation using Sephadex LH-20 column eluted with MeOH and subsequent TLC analysis, seven fractions were obtained (A–G). Fraction 5B was further fractionated using a linear gradient from 20% to 50% MeCN/H_2_O over 20 min by an Agilent Infinity 1290 HPLC-DAD (Agilent Technologies) system using a phenyl-hexyl column (Luna, 100 Å, 250 × 10 mm, i.d., 5 μm, Phenomenex), with a flow rate of 4 mL/min. Each fraction was again loaded on HPLC-DAD for the last purification step to afford compound **5** (lysolipin M; 2 mg).

#### Lysolipin I (1)

Yellow crystal; 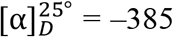 (c 1.5, CH_3_OH); UV(CH_3_CN/H_2_O) λ_max_ 245, 275, 310, 350 and 405 nm; ECD *λ*_ext_ (*Δ*_*ε*_) (CH_3_OH) 218 (+11), 314 (+3), 241 (–40), 278 (+22), 332 (– 5) nm ; ^1^H and ^13^C NMR data, see **Table 1**. HRESIMS m/z 598.1119 [M + H]^+^ (calcd 598.1111 for C_29_H_24_NO_11_Cl, Δ 1.34 ppm).

#### Lysolipin J (2)

Yellow crystal; 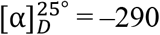 (c 1, CH_3_OH); UV(CH_3_CN/H_2_O) λ_max_ 245, 275, 310, 350 and 405 nm; ^1^H and ^13^C NMR data, see **Table 1**. HRESIMS m/z 584.0964 [M + H]^+^ (calcd 584.0954 for C_28_H_22_NO_11_Cl, Δ 1.7 ppm).

#### Lysolipin K (3)

Yellow crystal; 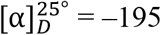 (c 1.5, CH_3_OH); UV(CH_3_CN/H_2_O) λ_max_ 245, 275, 310, 350 and 405 nm; ^1^H and ^13^C NMR data, see **Table 1**. HRESIMS m/z 584.0964 [M + H]^+^ (calcd 584.0954 for C_28_ H_22_NO_11_Cl, Δ 1.7 ppm).

#### Lysolipin L (4)

Yellow crystal; 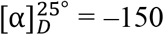 (c 1.5, CH_3_OH); UV(CH_3_CN/H_2_O) λ_max_ 270, 340 and 400 nm; ECD *λ*_ext_ (*Δ*ε) (CH_3_OH) 220 (+5), 300 (+4), 235 (–10) nm, ^1^H and ^13^C NMR data, see

**Table 1.** HRESIMS m/z 612.0914 [M + H]^+^ (calcd 612.0903 for C_29_H_22_ClNO_12_, Δ 1.8 ppm).

#### Lysolipin M (5)

Yellow crystal; 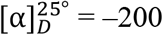 (c 1.5, CH_3_OH); UV(CH_3_CN/H_2_O) λ_max_ 230, 255, 320, 370 and 410 nm; ECD *λ*_ext_ (*Δ*ε) (CH_3_OH) 220 (+4), 300 (+3), 235 (–9) nm, ^1^H and ^13^C NMR data, see **Table 2**. HRESIMS m/z 700.1426 [M + H]^+^ (calcd 700.1428 for C_33_H_30_NO_14_Cl, Δ –0.3 ppm).

### Cytotoxicity Assay

Cell growth and cytotoxicity were assessed using the human epithelial prostate cancer cell line LNCaP (RRID:CVCL_0395) and its hormone-refractory derivative C4-2B (RRID:CVCL_4784), as well as two derivative drug-resistant sublines (C4-2BR and LNCaPR, respectively),^30^ all of which carry wild-type *RAS* alleles. All cell lines were cultured and maintained in RPMI-1640 medium containing glutaMAX™-I (Gibco, Invitrogen, Carlsbad, CA, United States), supplemented with 10% fetal bovine serum (FBS). Cells were seeded into 6-well plates at 3 × 10^5^ cells/well. After cells are attached, the medium in each well was replaced with 2 mL of fresh warm medium containing 10μM of the different compounds or a vehicle. Cell proliferation dynamics were monitored in real-time using a lens-free Cellwatcher microscopy device (PHIO, Germany). The cell growth curves were generated with the analysis module available from PHIO to determine the total area covered by cells. Cytotoxicity was measured as an endpoint assay of drug-exposure using the CellTox Green Cytotoxicity Assay kit according to manufacturer’s instructions. Briefly, CellTox green cytotoxicity reagent was added to the media at a final concentration of 1X, and the relative cytotoxicity was calculated relative to the control well treated with vehicle.

### Molecular Docking Studies

All the crystal structures were fetched from the PDB site, and the grid box parameters were manually adjusted to ensure complete encapsulation of the Topoisomerase IIβ active site, including the DNA cleavage region and adjacent residues involved in ligand binding. The grid dimensions were selected to fully accommodate the ligand and allow sufficient conformational flexibility during docking while preventing non-specific binding outside the catalytic pocket. The protein preparation was done by VEGA ZZ 3.1.1.4.^31^ Lysolipin I and the native ligands were drawn in 2D and non-minimized 3D forms using ChemBioDraw Ultra 14.0 and Chem-Bio3D Ultra 14.0, respectively. The lowest-energy conformers were obtained by semi-empirical energy minimisation using the MOPAC algorithm.by the MOPAC algorithm of energy minimization. The docking simulation was performed by Autodock Vina and MOE after writing the file formats in PDBQT using AutoDock Tools 1.5.7.^32^ All docking calculations were conducted using the default parameters implemented in each software package, including grid generation and scoring functions, with no additional parameter optimization, in order to ensure reproducibility of the docking experiments. Visualization and exporting images were accomplished by Chimera software.^33^

### Antimicrobial Activity Test

The antimicrobial activity of compounds 1–5 was evaluated against the Gram-positive bacterium *Staphylococcus aureus* strain 8325 and the fungal pathogen *Aspergillus flavus* IBT30114. *S. aureus* was cultured in Mueller–Hinton Broth (MHB), while *A. flavus* was maintained on Potato Dextrose Agar (PDA) at 30 °C for 7 days to ensure adequate sporulation prior to testing.

Minimum inhibitory concentrations (MICs) were determined using the broth microdilution method. Two-fold serial dilutions (64–0.03125 µg/mL) were prepared in MHB for *S. aureus* and Potato Dextrose Broth (PDB) for *A. flavus* in 96-well microplates. Each well was inoculated to a final concentration of 5×10^5^ CFU/mL for *S. aureus* and 1×10^5^ spores/mL for *A. flavus*.

The plates were incubated at 37 °C for 24 h (*S. aureus*) and 30 °C for 48 h (*A. flavus*). MIC values were defined as the lowest concentration of compound that completely inhibited visible growth. All experiments were performed in triplicate (*n* = 3).^34,35^

## Data Availability

The NMR data for compounds **1–5** have been deposited in the Natural Products Magnetic Resonance Database (NP-MRD; www.np-mrd.org) with the accession numbers for lysolipin J (NP0354059), lysolipin K (NP0354060), lysolipin L (NP0354061), and lysolipin M (NP0354062). The MS data are deposited at GNPS with the number of ID=d7aa0eb9e3ea470088dd28a50f0c39ea Genomic information is available in NCBI under GenBank PRJNA1359755 (*Streptomyces* sp. P8-2B18).

## Supporting Information

The Supporting Information is available free of charge at the ACS Publications website.

Co-cultivation of *Streptomyces* sp. P8-2B18 with fungal pathogens; 1D and 2D NMR, UV, HRESIMS data of compounds **1–4**; BGC annotation of *Streptomyces* sp. P8-2B18; phylogenetic tree of *Streptomyces* sp. P8-2B18 and other bacteria (PDF).

## AUTHOR INFORMATION

### Notes

The authors declare no competing financial interest.

## ACKNOWLEDGMENT

This study was supported by Egyptian governmental scholarship. Thanks to the support from AJC, Andersen and U. Rubeziute, DTU Metabolomics Core; The NMR Center DTU and the Villum Foundation are acknowledged for access to the 800 MHz NMR spectrometer. Thanks to support from Carlsberg Infrastructure (CF20-0177). K.Y.L. thanks to the Novo Nordisk Foundation (NNF22OC0079187). Y.W.L and L.D. thanks to Novo Nordisk Foundation (NNF23OC0082881 and NNF23OC0082882). L.D and M. L thank Danish National Research Foundation (DNRF137) as part of the Center for Microbial Secondary Metabolites (CeMiSt).

