## Supplemental Data 1 for "Antimicrobial and Cytotoxic Lysolipins I–M Isolated from *Streptomyces* sp. P8-2B18"

**Table S6.** Annotation of biosynthetic genes of lysolipins in *Streptomyces* sp. P8-2B18………..36

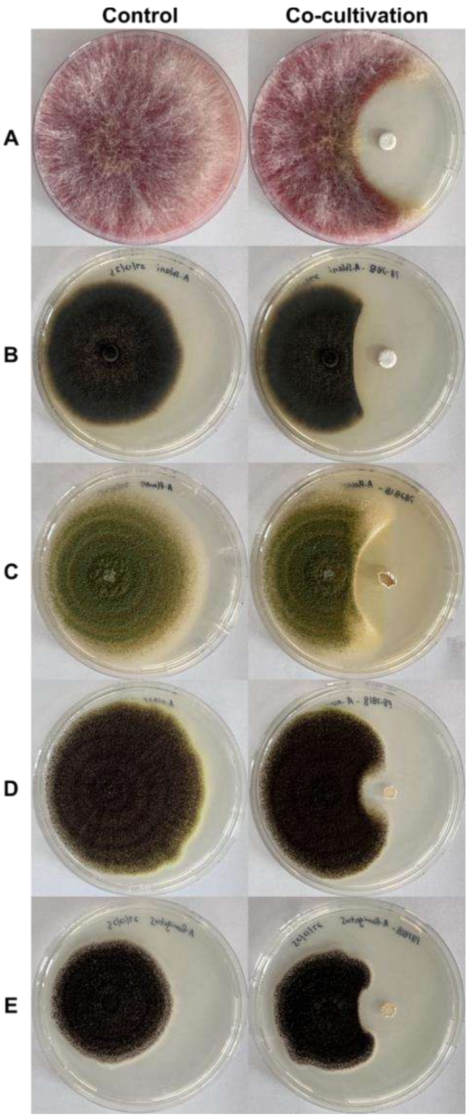

**Figure S1.** Co-cultivation of *Streptomyces* sp. P8-2B18 with fungal pathogens A) *Fusarium graminearum* IBT42824, B) *Alternaria solani*, C) *Aspergillus flavus*, D) *Aspergillus niger*, and E) *Aspergillus fumigatus* on PDA plates.

**Commented [LD1]:** @Kah Yean Lum • For the co-cultivation antifungal plates, consider adding scale bars and briefly describing assay conditions (medium, inoculum type, incubation time) either in the figure caption or a short methods section.

**Commented [LD2R1]:** @Kah Yean Lum

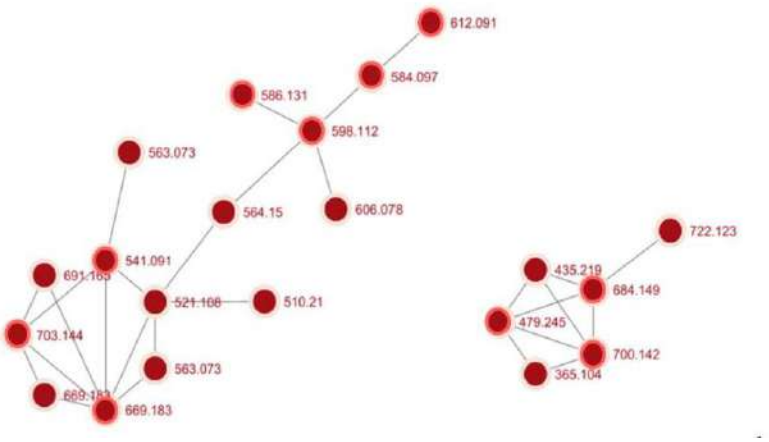

**Figure S2.** Molecular network cluster of lysolipin I (m/z 598.1111 $[M+H]^+$) and its derivatives

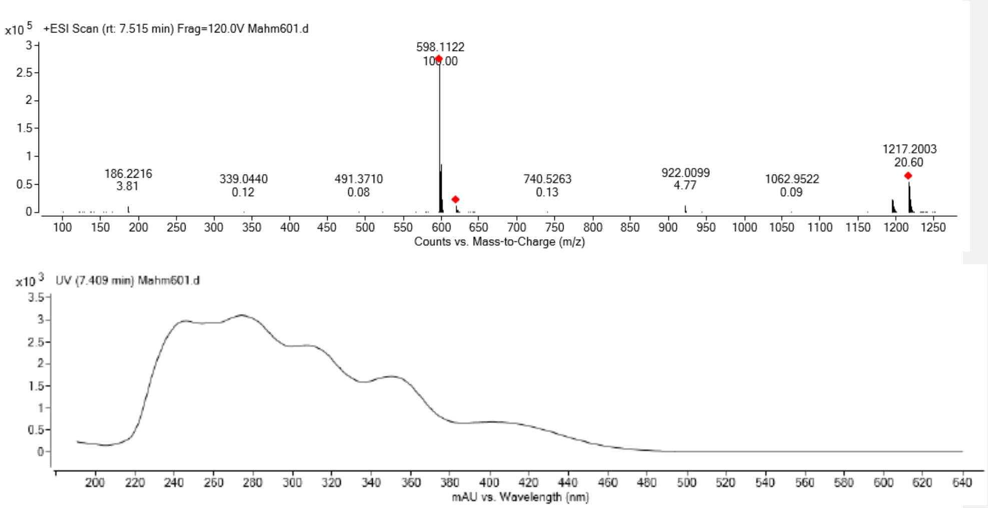

**Figure S3.** (+)-HRESIMS and UV spectra of lysolipin I (**1**)

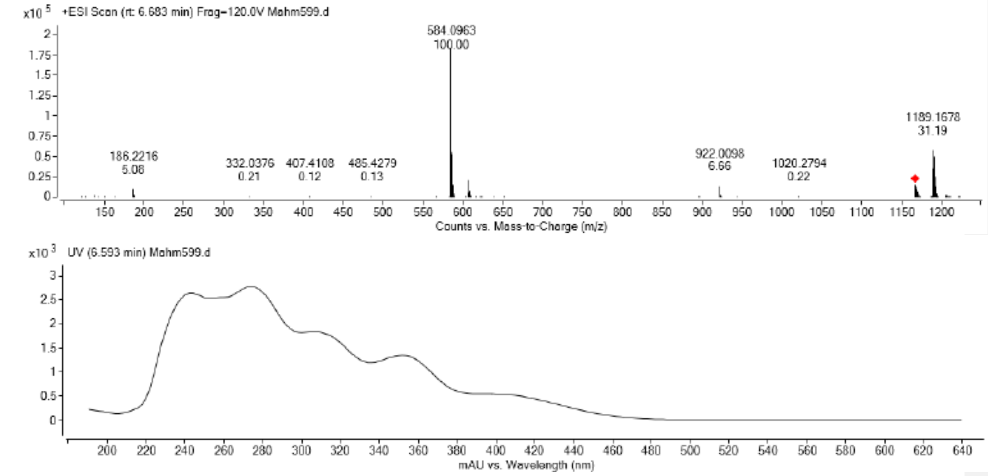

**Figure S4.** (+)-HRESIMS and UV spectra of lysolipin J (**2**)

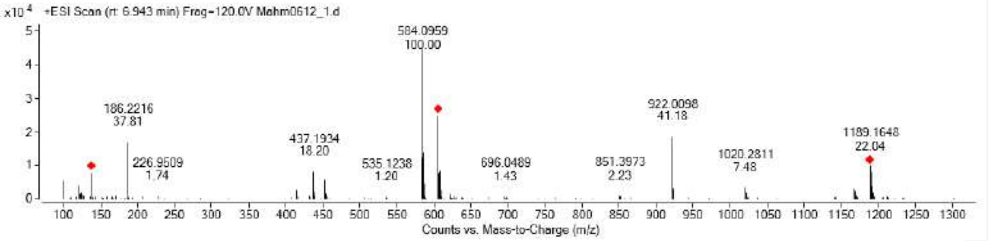

.

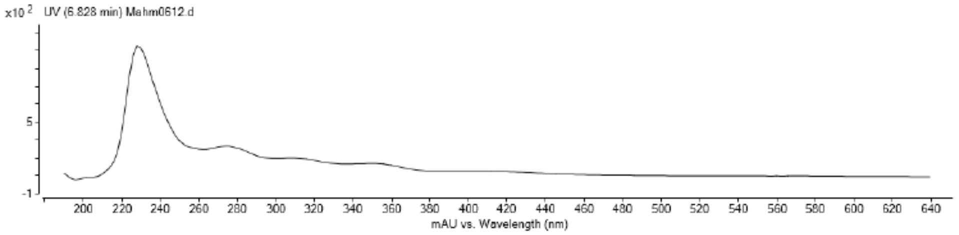

**Figure S5.** (+)-HRESIMS and UV spectra of lysolipin K(**3**).

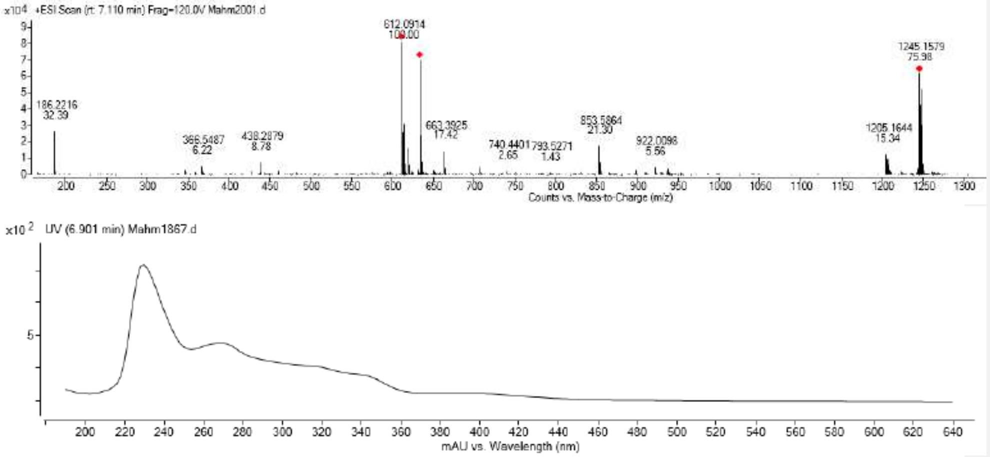

**Figure S6.** (+)-HRESIMS and UV spectra of lysolipin L (**4**).

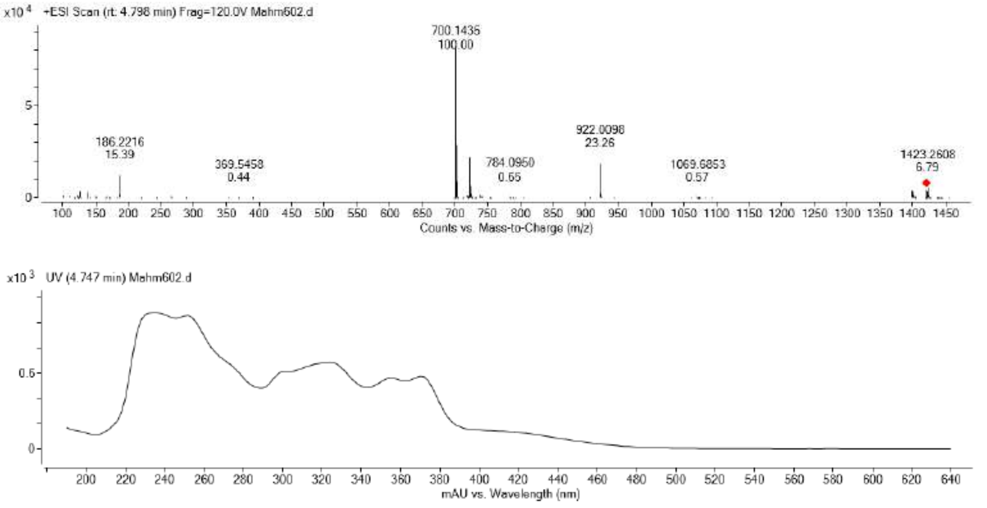

**Figure S7.** (+)-HRESIMS and UV spectra of lysolipin M (**5**).

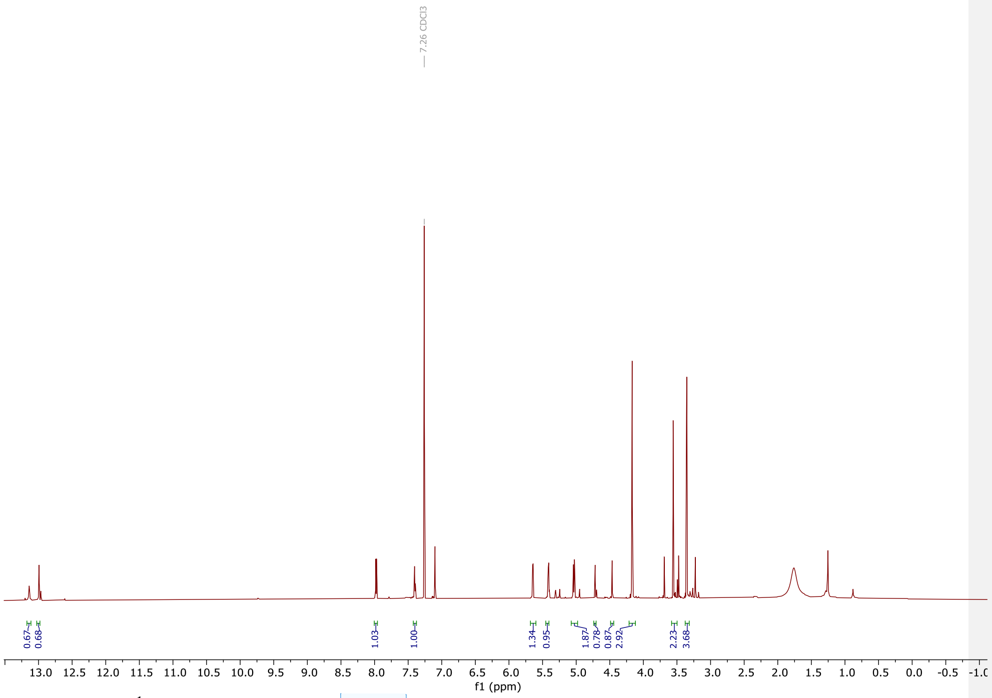

**Figure S8.** $^1$H NMR (800 MHz, $CDCl_3$) of lysolipin I (**1**)

**Commented [LD3]:** • Some SI figure captions are very brief. It would be useful if captions included the basic experimental details (e.g. “1H NMR (800 MHz, CDCl3) of lysolipin I (1)”) and the correct compound labels. Ensure that names and numbers are consistent between the main text and SI.
@Manar Magdy Mahmoud Mohamed

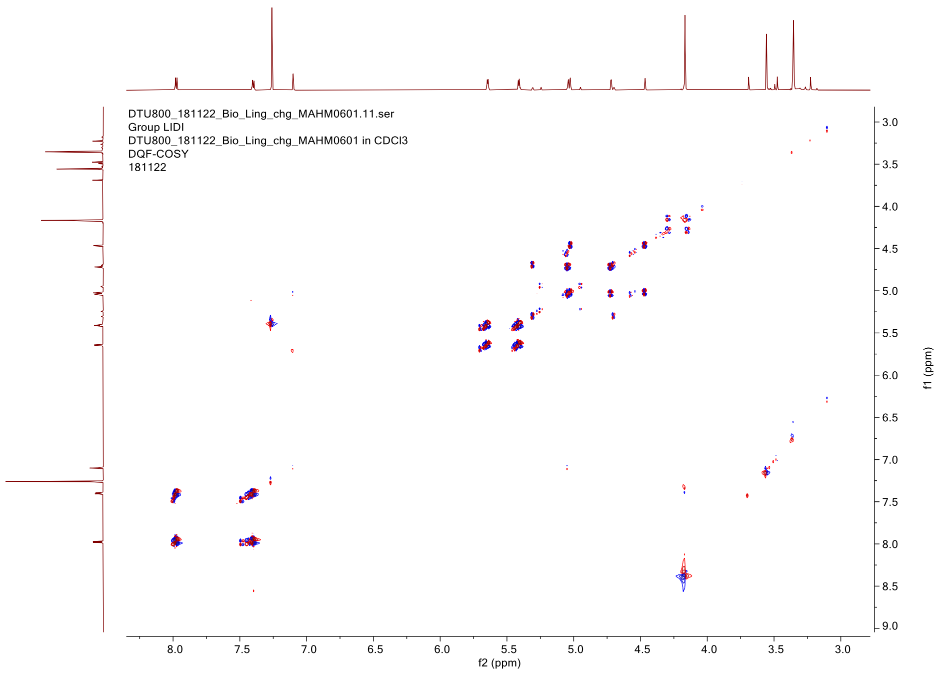

**Figure S9.** DQF-COSY (800 MHz, $CDCl_3$) of lysolipin I (**1**)

Commented [LD4]: @Manar Magdy Mahmoud Mohamed Please add MHZ for all the NMR spectra.

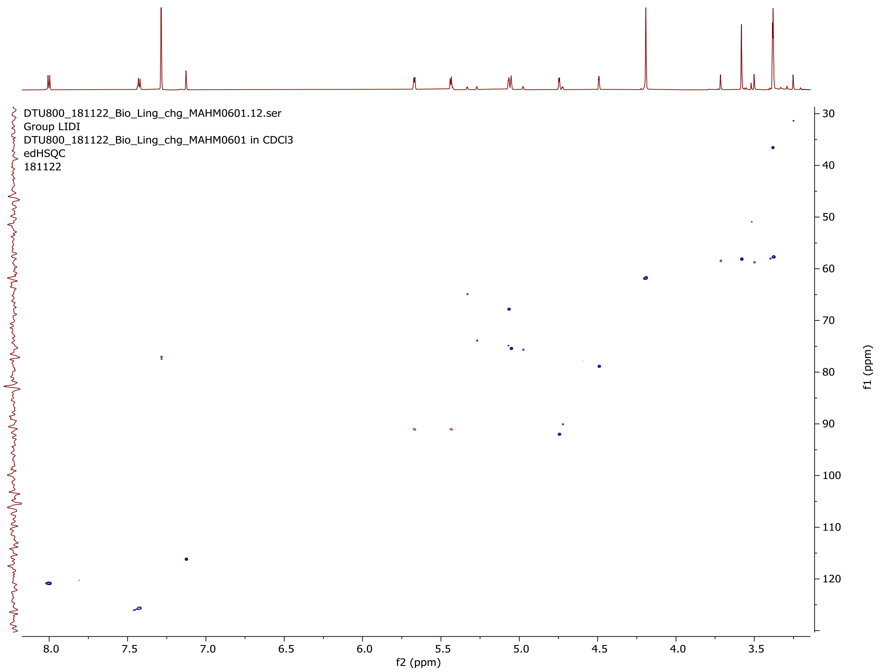

**Figure S10.** HSQC (200 MHz, $CDCl_3$) of lysolipin I (**1**)

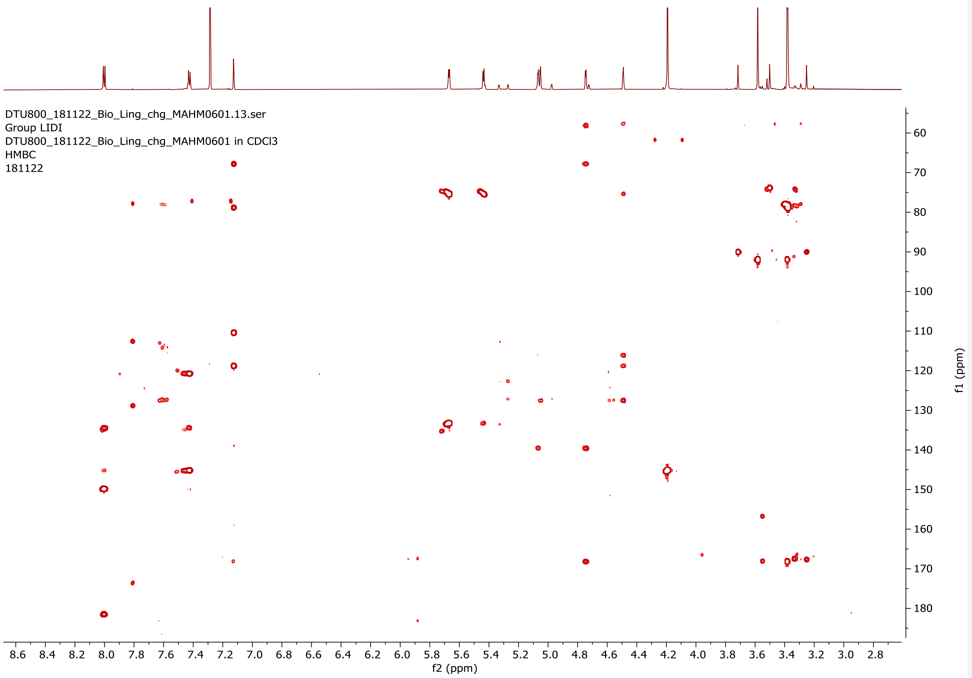

**Figure S11.** HMBC (200 MHz, $CDCl_3$) of lysolipin I (**1**).

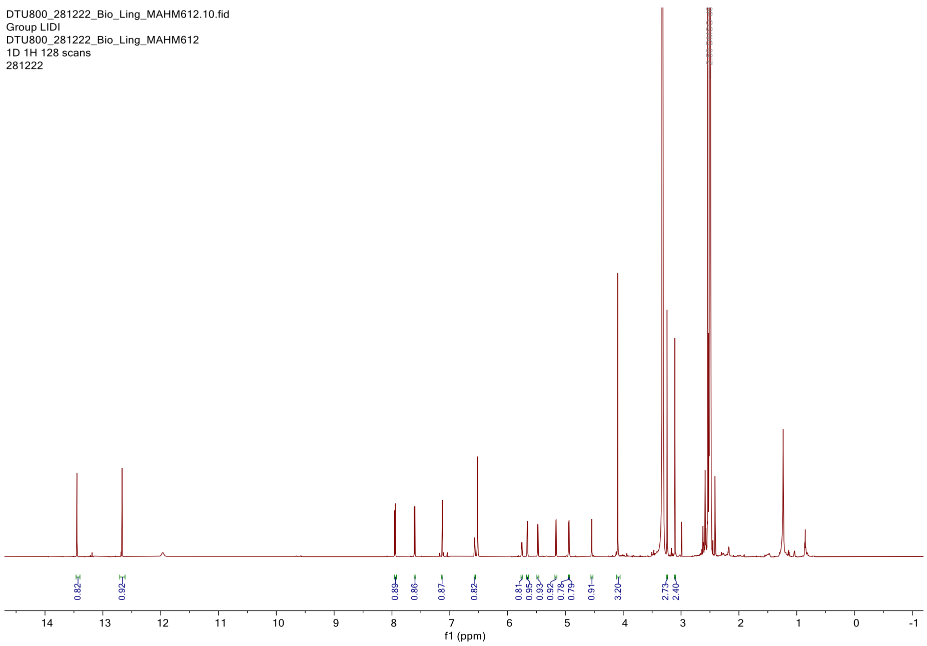

**Figure S12.** $^{1}H$ NMR (800 MHz, DMSO-$d_6$) of lysolipin J (**2**).

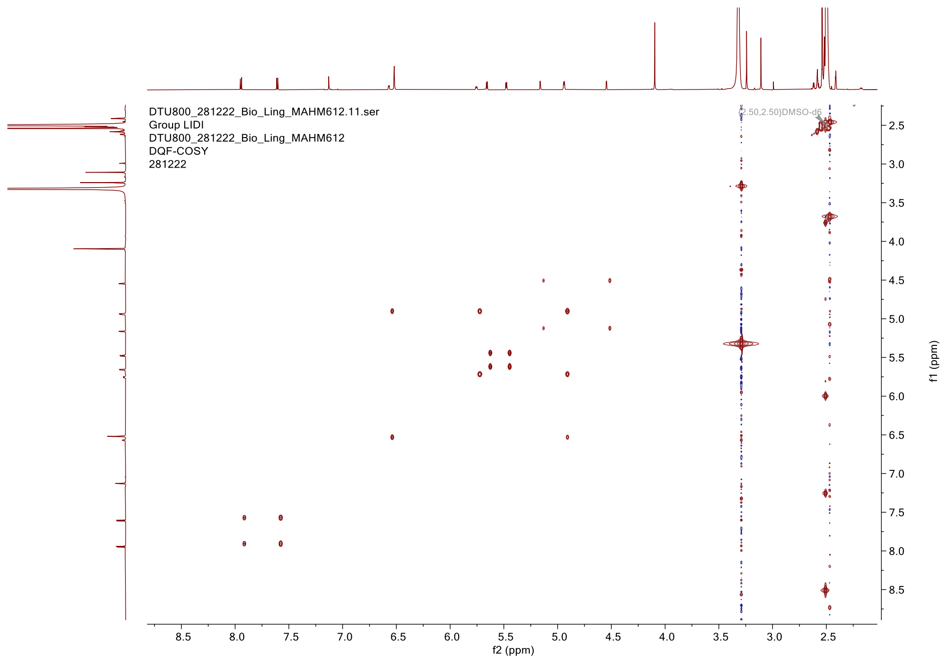

**Figure S13.** DQF-COSY (800 MHz, DMSO-*d*$_6$) of lysolipin J (**2**).

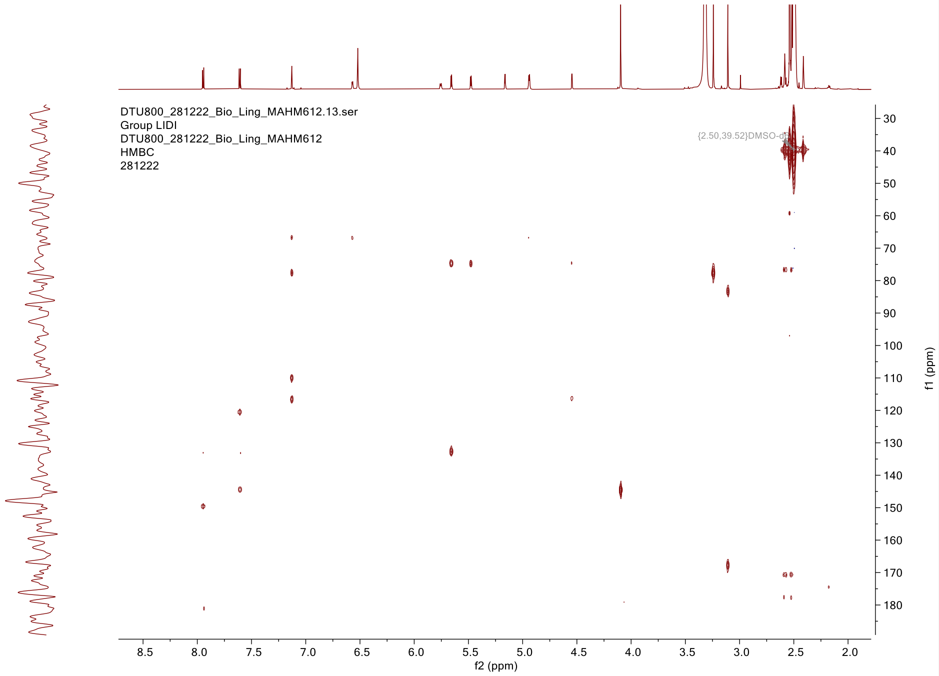

**Figure S14.** HMBC (200 MHz, DMSO-$d_6$) of lysolipin J (**2**).

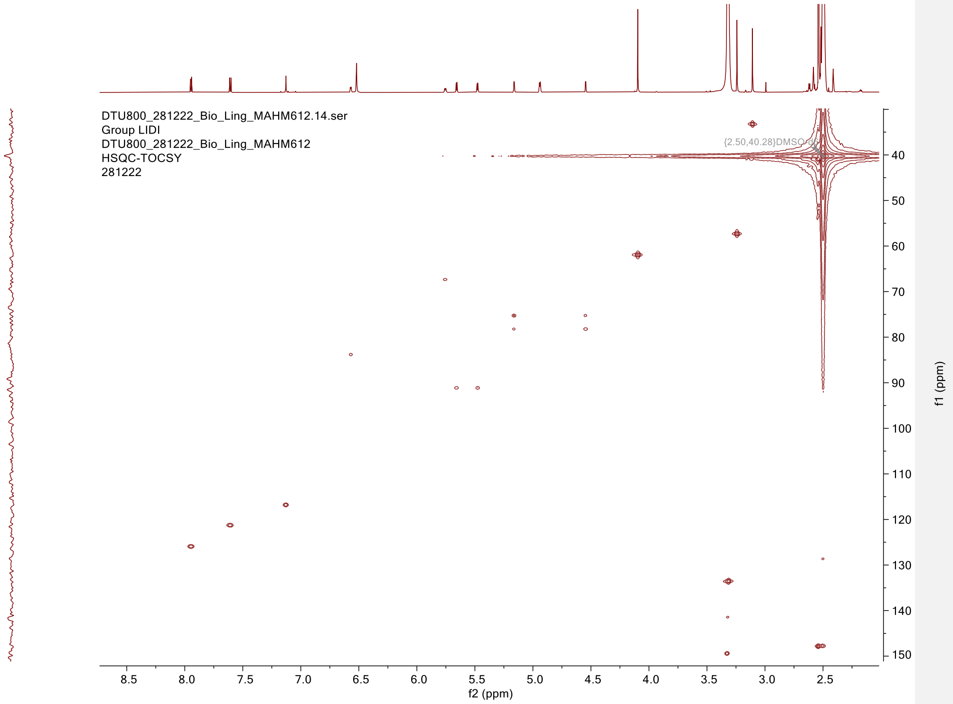

**Figure S15.** HSQC-TOCSY(200 MHz, DMSO-$d_6$) of lysolipin J (**2**).

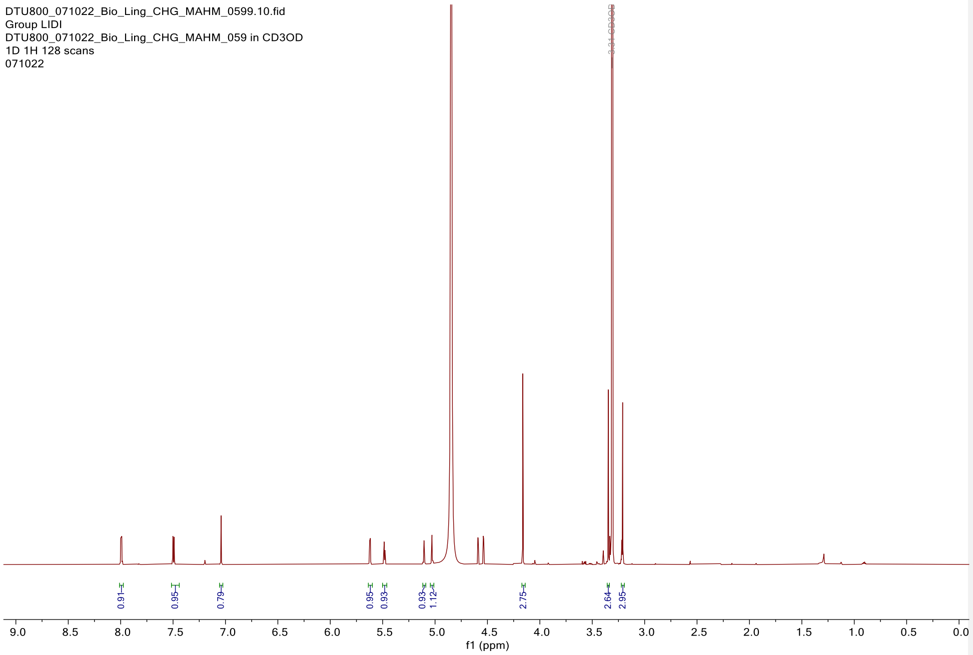

**Figure S16.** $^{1}H$ NMR (800 MHz, $CD_3OD$) of lysolipin K (**3**).

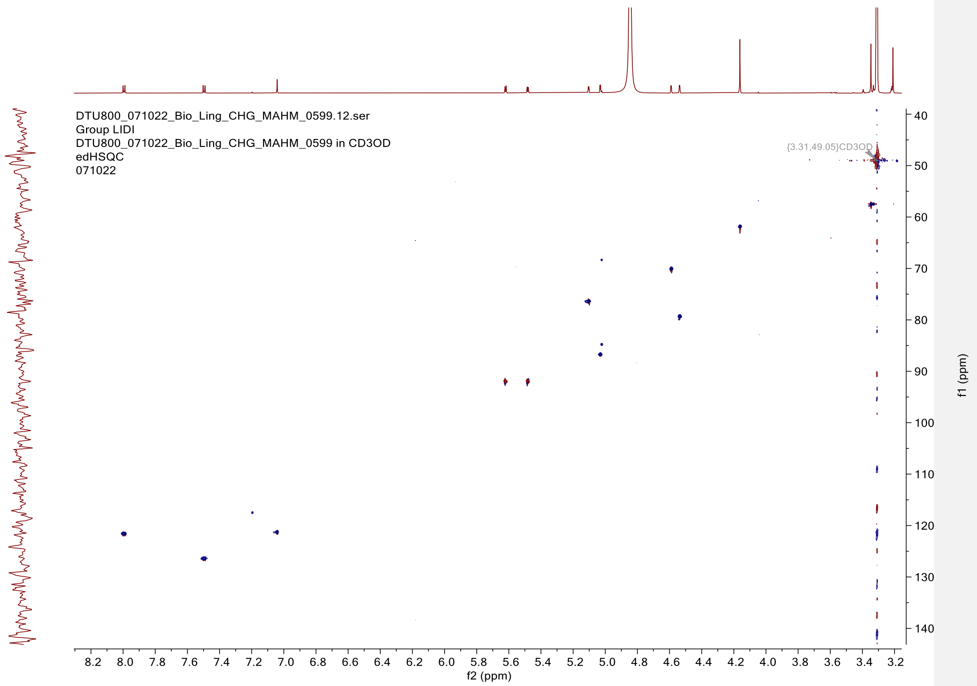

**Figure S17.** HSQC (200 MHz, $CD_3OD$) of lysolipin K (**3**).

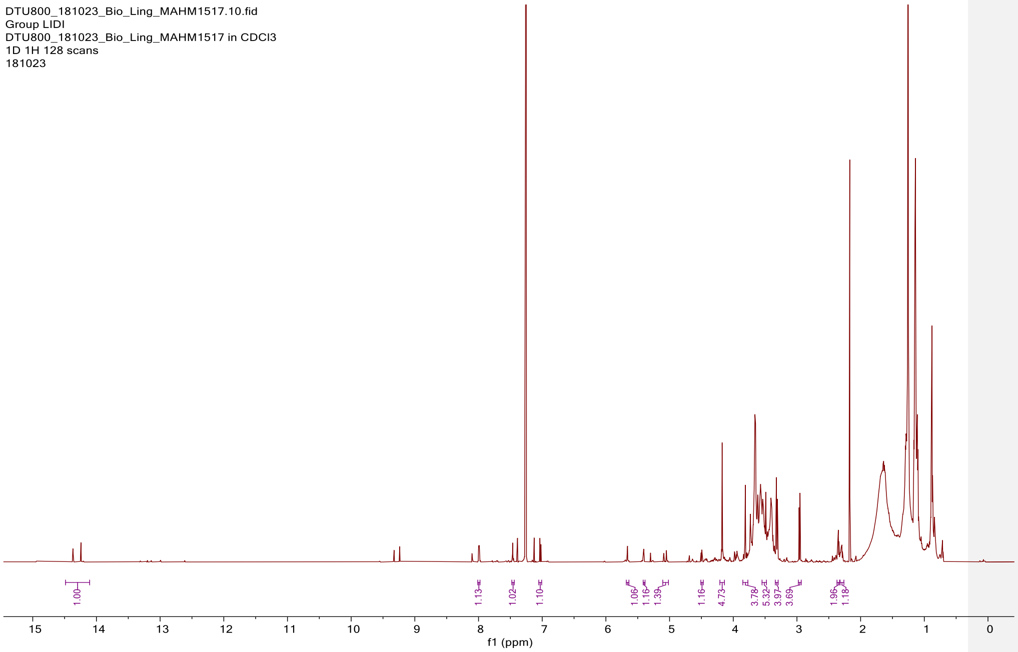

**Figure S18.** [1]H NMR (800 MHz, $CDCl_3$) of lysolipin L (**4**).

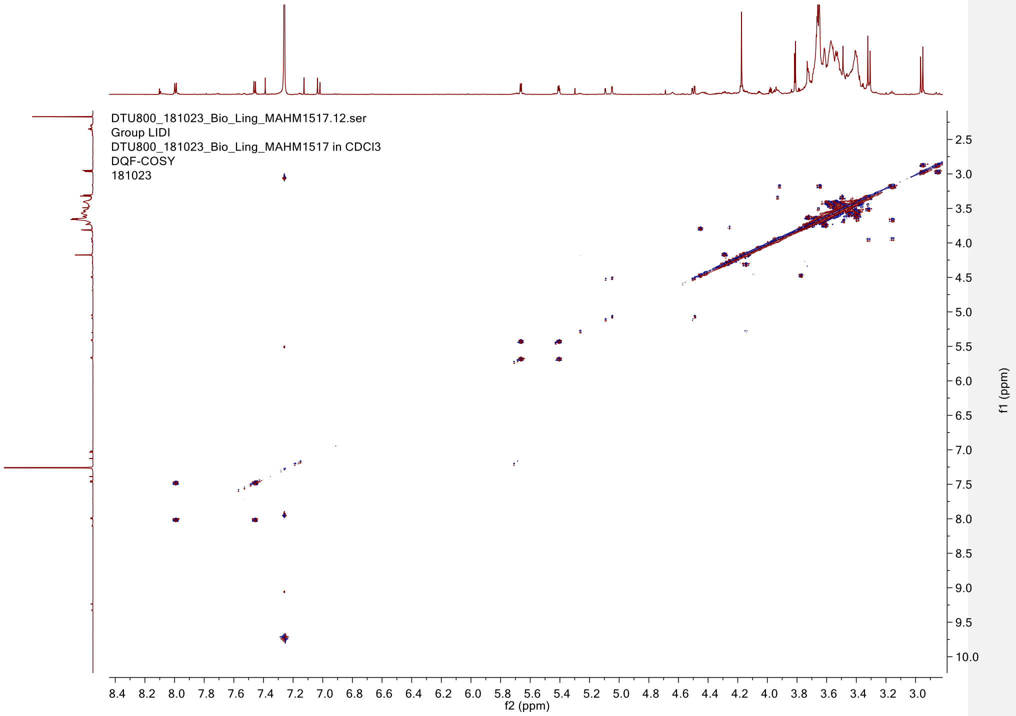

**Figure S19.** DQF-COSY (800 MHz, $CDCl_3$) of lysolipin L (**4**).

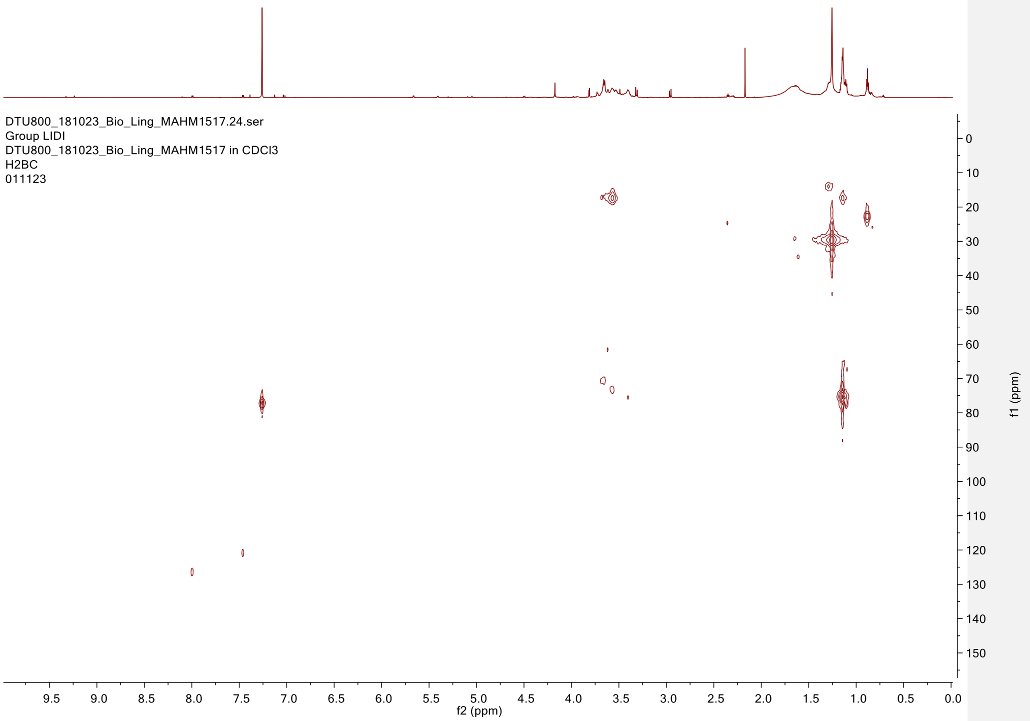

**Figure S20.** H2BC (200 MHz, $CDCl_3$) of lysolipin L (**4**).

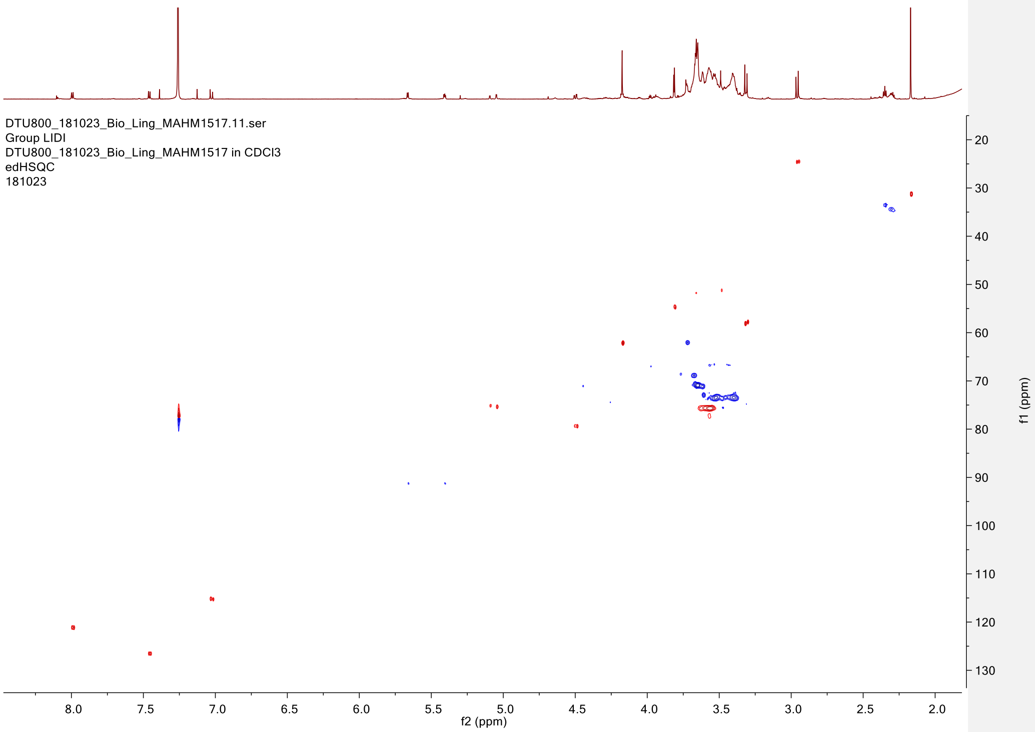

**Figure S21.** HSQC (200 MHz, $CDCl_3$) of lysolipin L (**4**).

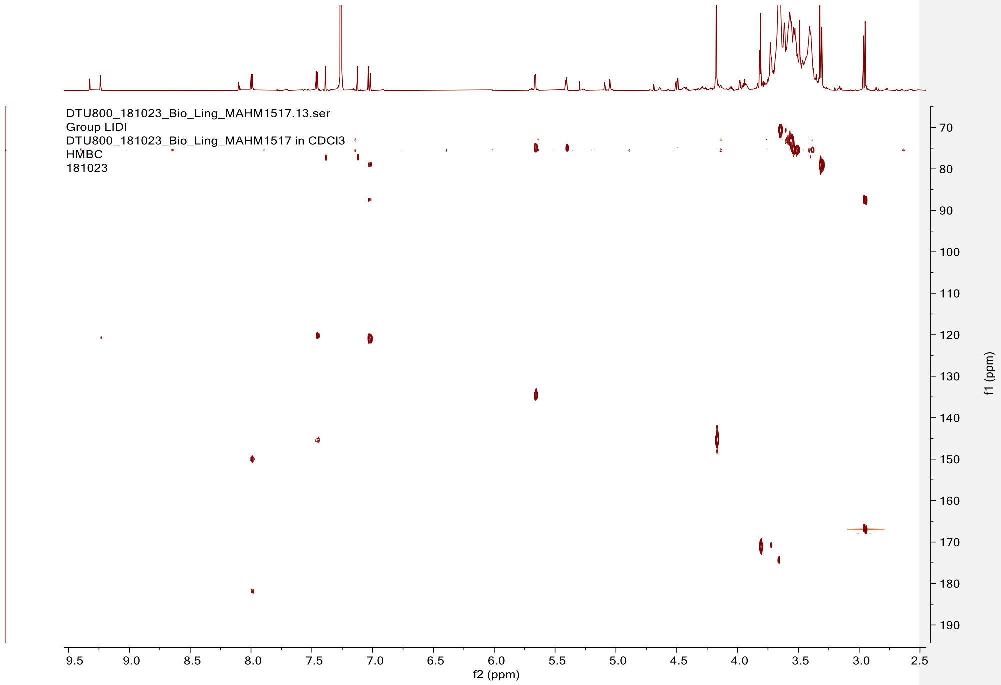

**Figure S22.** HMBC (200 MHz, $CDCl_3$) of lysolipin L (**4**).

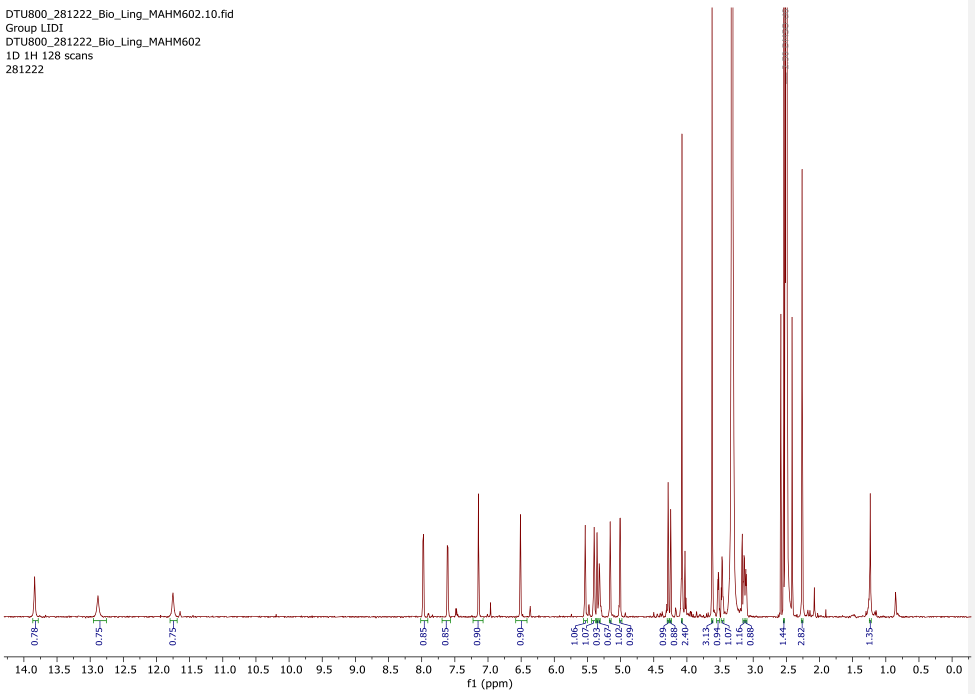

**Figure S23.** $^{1}$H NMR (800 MHz, DMSO-$d_6$) of lysolipin M (**5**).

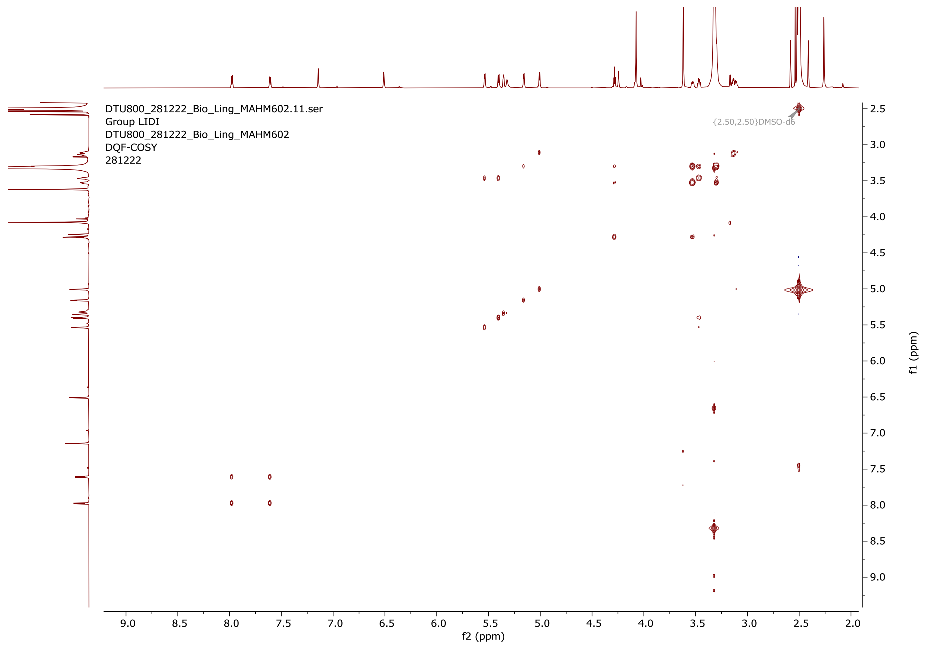

**Figure S24.** DQF-COSY (800 MHz, DMSO-$d_6$) of lysolipin M (**5**).

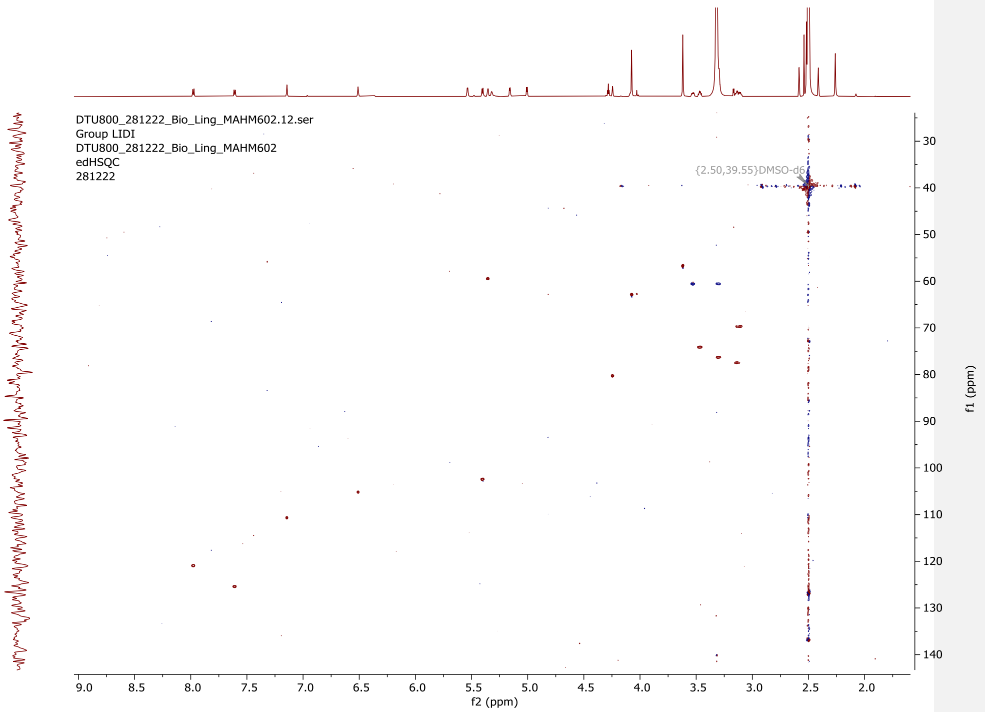

**Figure S25.** HSQC (200 MHz, DMSO-$d_6$) of lysolipin M (**5**).

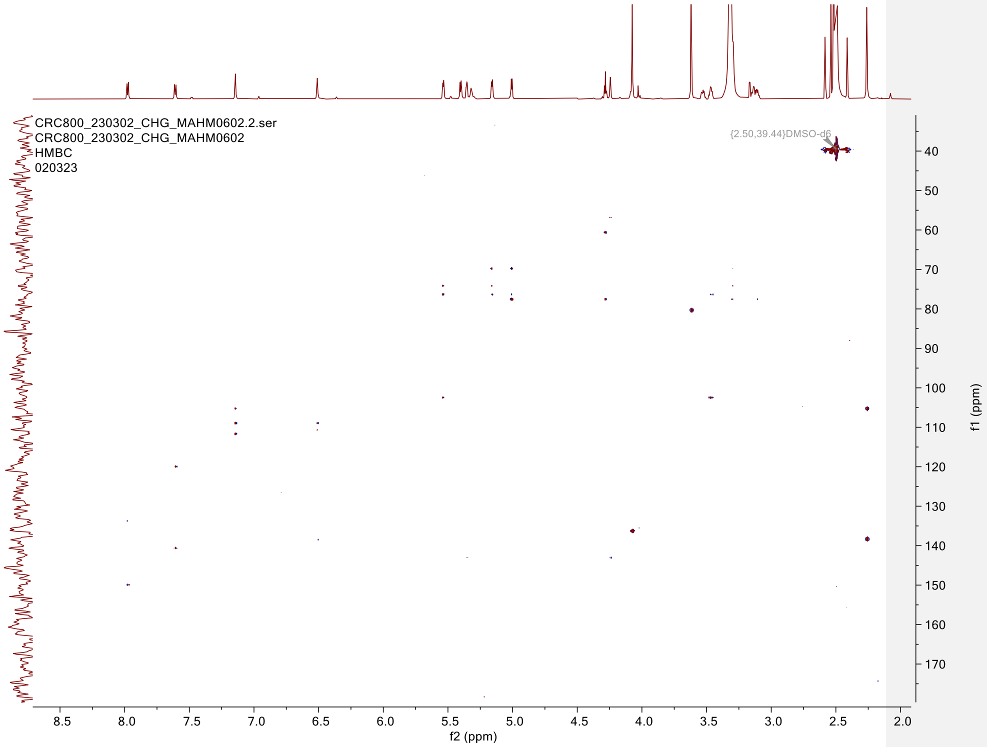

**Figure S26.** HMBC (200 MHz, DMSO-$d_6$) of lysolipin M (**5**).

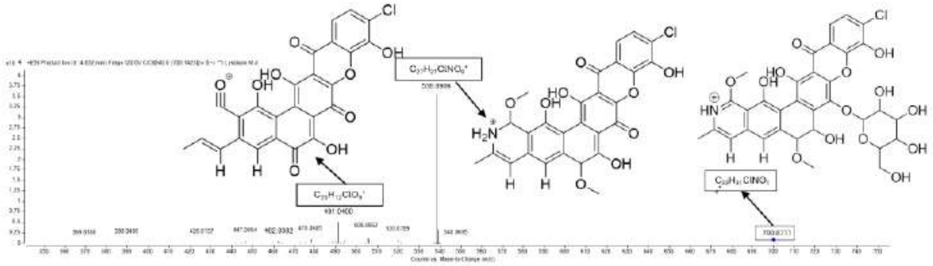

**Figure S27.** (+)-HRESIMS/MS (40 eV) spectrum of lysolipins M (**5**).

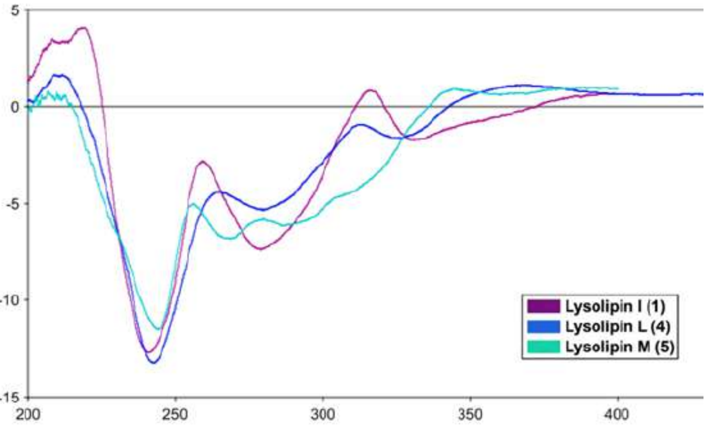

**Figure S28.** Electronic Circular Dichroism (ECD) spectra of lysolipins (**1**, **4** and **5**).

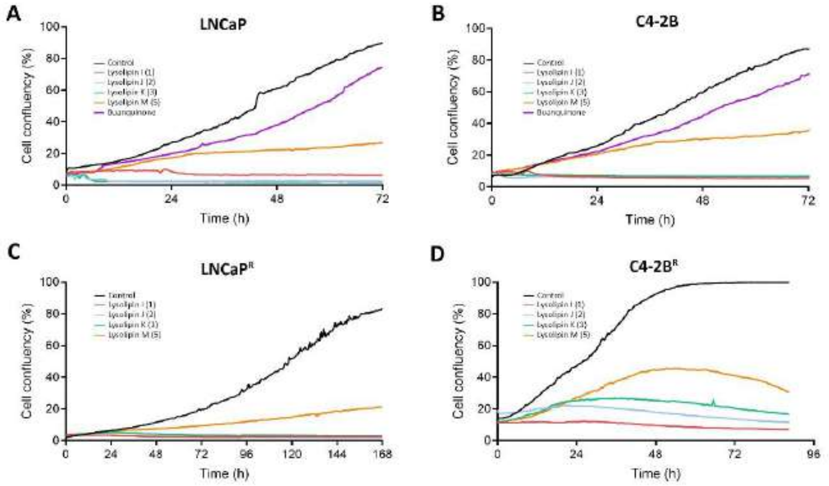

**Figure S29.** Effect of lysolipins on cell growth. Label-free confluence measurements of (A) LNCaP, (B) C4-2B, (C) LNCaPR, and (D) C4-2BR prostate cancer cells, showing high growth-inhibitory effects for lysolipins (**1**-**3**) and moderate effects for buanquinone.

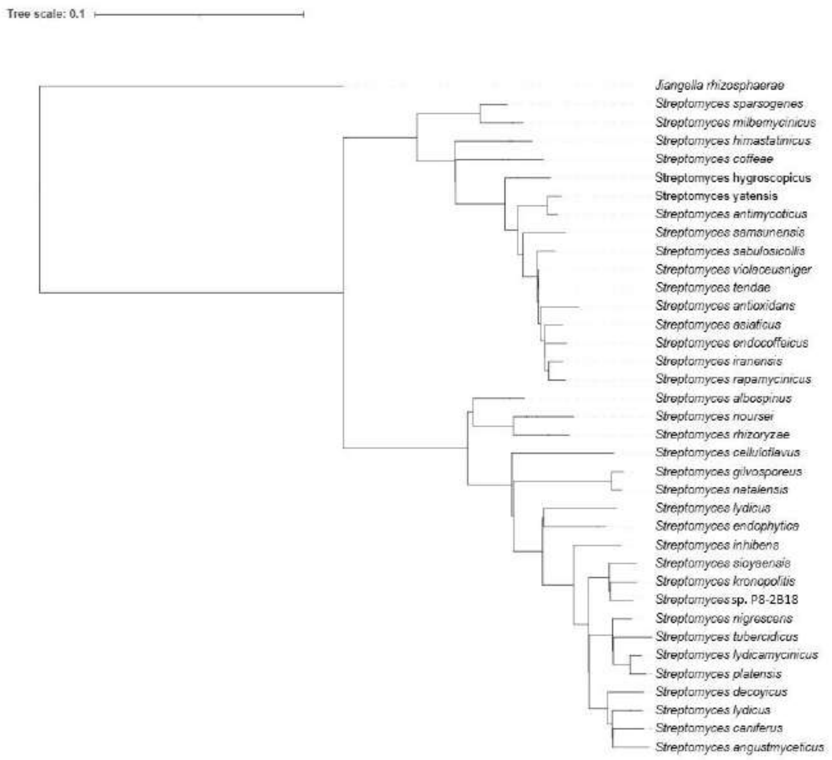

**Figure S30.** Phylogenetic tree of *Streptomyces* sp. P8-2B18 and other related *Streptomyces* spp.

**Table S12.** $^1$H (800 MHz) and $^{13}$C (200 MHz) NMR Data, along with HMBC and COSY correlations, for **l**ysolipin I (**1**) in DMSO-$d_6$.

| pos. | **1** in $CDCl_3$ | | | |
|---|---|---|---|---|
| | δ$_C$, type | δ$_H$ (*J* in Hz) | COSY | HMBC |
| **1** | 134.8, C | | — | — |
| **2** | 125.8, CH | 7.40 (d, 8.6) | H-3 | C-4, C-6 |
| **3** | 120.8, CH | 7.97 (d, 8.6) | H-2 | C-1, C-5, C-10 |
| **4** | 121.0, C | | — | — |
| **5** | 150.1, C | | — | — |
| **6** | 145.5, C | | — | — |
| **8** | -[a] | | — | — |
| **9** | 109.1, C | | — | — |
| **10** | 181.9, C | | — | — |
| **11** | 152.0, C | | — | — |
| **12** | 111.3, C | | — | — |
| **13** | 127.6, C | | — | — |
| **14** | 133.6, C | | — | — |
| **15** | 75.2, CH | 5.02 (d, 3.0) | H-16 | — |
| **16** | 78.9, CH | 4.47 (d, 2.9) | H-15 | C-18, C-22 |
| **17** | -[a] | | — | — |
| **18** | 119.5, C | | — | — |
| **19** | 153.2, C | | — | C-24, C-26 |
| **20** | 109.4, C | | — | — |
| **21** | 140.0, C | | — | — |
| **22** | 116.2, CH | 7.10 (s) | | C-16, C-18, C-20, C-23 |
| **23** | 67.5, CH | 5.04, (d, 3.9) | H-24 | C-21 |
| **24** | 92.0, CH | 4.72, (d, 3.8) | H-23 | C-21, C-26, C-36, C-37 |
| **26** | 168.5, C | | | |
| **28** | 91.1, $CH_2$ | 5.41 (d, 5.7), 5.64 (d, 5.7) | | C-14, C-15 |
| **31** | 61.9, $CH_3$ | 4.17 (s) | | C-6 |
| **33** | 58.0, $CH_3$ | 3.56 (s) | | C-16 |
| **36** | 57.6, $CH_3$ | 3.35 (s) | | C-24 |
| **37** | 36.6, $CH_3$ | 3.36 (s) | | C-24, C-26 |
| **38** | | 13.98 (s) | | |
| **39** | | 13.19 (s) | | |

**Table S23**. $^{1}$H (800 MHz) and $^{13}$C (200 MHz) NMR Data, along with HMBC and COSY correlations, for lysolipin J (**2**) in DMSO-$d_6$.

| pos. | **2** in DMSO-$d_6$ | | | |
| --- | --- | --- | --- | --- |
| | $\delta_C$, type | $\delta_H$ (*J* in Hz) | COSY | HMBC |
| **1** | 133.1, C | | — | — |
| **2** | 125.2, CH | 7.60 (d,8.7) | H-3 | C-4, C-6 |
| **3** | 120.1, CH | 7.94 (d, 8.6) | H-2 | C-1, C-5, C-10 |
| **4** | 120.5, C | | — | — |
| **5** | 149.6, C | | — | — |
| **6** | 144.4, C | | — | — |
| **8** | -[a] | | — | — |
| **9** | 107.8, C | | — | — |
| **10** | 181.1, C | | — | — |
| **11** | 150.2, C | | — | — |
| **12** | 111.2, C | | — | — |
| **13** | 128.4, C | | — | — |
| **14** | 132.3, C | | — | — |
| **15** | 74.5, CH | 5.16 (d, 2.7) | H-16 | — |
| **16** | 77.5, CH | 4.55 (d, 2.7) | H-15 | C-15, C-22 |
| **17** | -[a] | | — | — |
| **18** | 116.8, C | | — | — |
| **19** | 157.7, C | | — | — |
| **20** | 109.8, C | | — | — |
| **21** | -[a] | | — | C-16, C-18, C-20, C-23 |
| **22** | 116.1, CH | 7.04 (s) | H-24 | |
| **23** | 66.6, CH | 4.95 (d, 3.1) | H-23 | |
| **24** | 83.1, CH | 4.94 (d, 3.0) | — | — |
| **26** | 167.8, C | | — | C-14, C-15 |
| **28** | 92.0, $CH_2$ | 5.48 (d, 5.7), 5.66 (d, 5.7) | — | C-6 |
| **31** | 61.2, $CH_3$ | 4.10 (s) | — | C-16 |
| **33** | 56.6, $CH_3$ | 3.24 (s) | — | — |
| **36** | | | — | C-24, C-26 |
| **37** | 32.5, $CH_3$ | 3.11 (s) | — | — |
| **38** | | 13.45 (s) | — | C-18, C-19, C-20 |
| **39** | | 12.67 (s) | — | C-9, C-11, C-12 |

**Table S34**. $^{1}H$ (800 MHz) and $^{13}C$ (200 MHz) NMR Data, along with HMBC and COSY correlations, for lysolipin L (**4**) in $CDCl_3$

| pos. | **4** in $CDCl_3$ | | | |
|---|---|---|---|---|
| | $\delta_C$, type | $\delta_H$ ($J$ in Hz) | COSY | HMBC |
| **1** | 135.4, C | | — | — |
| **2** | 126.1, CH | 7.46 (d, 8.64) | H-3 | C-4, C-6 |
| **3** | 120.7, CH | 7.99 (d, 8.66) | H-2 | C-1, C-5, C-10 |
| **4** | 120.2, C | | — | — |
| **5** | 150.0, C | | — | — |
| **6** | 145.4, C | | — | — |
| **8** | -[a] | | — | — |
| **9** | -[a] | | — | — |
| **10** | 181.8, C | | — | — |
| **11** | -[a] | | — | — |
| **12** | -[a] | | — | — |
| **13** | -[a] | | — | — |
| **14** | 134.6, C | | — | — |
| **15** | 75.1, CH | 5.04 (d, 3.1) | H-16 | C-14, C-28 |
| **16** | 79.4, CH | 4.49 (d, 3.0) | H-15 | C-18, C-22, C-33 |
| **17** | -[a] | | — | — |
| **18** | 121.0, C | | — | — |
| **19** | -[a] | | — | — |
| **20** | -[a] | | — | — |
| **21** | -[a] | | — | — |
| **22** | 114.7, CH | 7.04 (s) | — | C-16, C-23 |
| **23** | 87.4, C | | — | — |
| **24** | 171.2, C | | — | — |
| **26** | 166.5, C | | — | — |
| **28** | 91.2, $CH_2$ | 5.41 (dd, 5.6, 3,8), 5.66 (d, 5.6) | — | C-14, C-15 |
| **31** | 62.1, $CH_3$ | 4.17 (s) | — | C-6 |
| **33** | 58.2, $CH_3$ | 3.32(s) | — | C-16 |
| **36** | 54.7, $CH_3$ | 3.81 (s) | — | C-24 |
| **37** | 24.5, $CH_3$ | 2.97 (s) | — | C-23, C-26 |
| **38** | | 14.2 (s) | — | — |
| **39** | | 14.4 (s) | — | — |

**Table S45.** $^{1}$H (800 MHz) and $^{13}$C (200 MHz) NMR Data, along with HMBC and COSY correlations, for lysolipin M (**5**) in DMSO-*d*$_6$.

| **5** in DMSO-d$_6$ | | | | | | | | | |
|---|---|---|---|---|---|---|---|---|---|
| pos. | δC, type | δH (*J* in Hz) | COSY | HMBC | pos. | δC, type | δH (*J* in Hz) | COSY | HMBC |
| **1** | 133.8, C | — | — | — | **26** | 136.6, C | | — | — |
| **2** | 125.3, CH | 7.61(d, 8.6) | C-3 | C-4, C-6 | **31** | | 11.75 (s) | — | — |
| **3** | 120.9, CH | 7.97 (d, 8.6) | C-2 | C-1, C-5 | **33** | 56.7, $CH_3$ | 3.62 (s) | — | C-16 |
| **4** | 120.1, C | — | — | — | **36** | 18.3, $CH_3$ | 2.26 (s) | — | C-23, C-24 |
| **5** | 150.0, C | — | — | — | **37** | 62.7, $CH_3$ | 4.08 (s) | — | C-26 |
| **6** | 140.6, C | — | — | — | **38** | | 12.9 (s) | — | — |
| **8** | -[a] | — | — | — | **39** | | 13.8 (s) | — | — |
| **9** | 108.8, C | — | — | — | **1'** | 102.4, CH | 5.41 (d, 7.6) | H-2' | C-15 |
| **10** | 186.5, C | — | — | — | **2'** | 74.0, CH | 3.47 (td, 8.2, 5.1) | H-1', H-3' | C-3' |
| **11** | 157.4, C | — | — | — | **2'-OH** | | 5.54 (d, 5.1) | — | — |
| **12** | 111.9, C | — | — | — | **3'** | 76.9, CH | 3.32 (m) | H-2', H-4' | C-4' |
| **13** | 112.8, C | — | — | — | **3'-OH** | | 5.16 (d, 5.0) | — | — |
| **14** | 143.2, C | — | — | — | **4'** | 70.6, CH | 3.11 (td, 8.8, 5.4) | H-3', H-5' | C-5' |
| **15** | 59.4, CH | 5.36 (d, 2.6) | C-16 | — | **4'-OH** | | 5.01 (d, 5.4) | — | — |
| **16** | 80.2, CH | 4.25 (brs) | C-15 | C-18 | **5'** | 77.4, CH | 3.14 (m) | H-4', H-6' | C-6' |
| **17** | -[a] | — | — | — | **6'** | 60.4, $CH_2$ | 3.53 (m); 3.30 (m) | H-5' | C-5' |
| **18** | 111.7, C | — | — | — | **6'-OH** | | 4.28 (t, 5.6) | — | — |
| **19** | 153.1, C | — | — | — | | | | | |
| **20** | 109.0, C | — | — | — | | | | | |
| **21** | 111.7, C | — | — | — | | | | | |
| **22** | 110.6, CH | 7.15 (s) | — | C-18, C-20, C-23 | | | | | |
| **23** | 105.1, CH | 6.51 (s) | — | C-20, C-22 | | | | | |
| **24** | 138.3, C | | — | — | | | | | |

**Table S5.** The BGC annotation of *Streptomyces* sp. P8-2B18.

| Region | Type | From | To | Similarity Confidence | Most similar known cluster | |
|---|---|---|---|---|---|---|
| Region 1.1 | T1PKS | 311,367 | 357,720 | | | |
| Region 1.2 | NRPS, NRPS-like, betalactone | 522,169 | 579,676 | Low | raimonol | terpene |
| Region 1.3 | hglE-KS, T1PKS | 847,664 | 899,265 | | | |
| Region 1.4 | ectoine | 1,279,295 | 1,289,711 | High | ectoine | other:ectoine |
| Region 1.5 | NI-siderophore | 1,372,363 | 1,402,168 | High | legonoxamine A/desferrioxamine B/legonoxamine B | other:other |
| Region 1.6 | NI-siderophore | 2,122,082 | 2,154,978 | Low | peucechelin | NRPS:Type I |
| Region 1.7 | crocagin, T2PKS | 2,274,091 | 2,358,213 | Medium | spore pigment | PKS |
| Region 1.8 | RiPP-like | 2,377,469 | 2,387,696 | | | |
| Region 1.9 | hydrogen-cyanide | 2,486,453 | 2,499,389 | Low | aborycin | ribosomal:RiPP |
| Region 1.10 | T3PKS | 2,529,608 | 2,570,672 | High | naringenin | PKS:Type III |
| Region 1.11 | RiPP-like | 2,590,600 | 2,602,528 | | | |
| Region 1.12 | NRPS, NRPS-like | 2,620,087 | 2,721,010 | Low | lipstatin | NRPS:Type I |
| Region 1.13 | azole-containing-RiPP, NAPAA, RRE-containing | 2,761,786 | 2,803,420 | Low | cyclothiazomycin | ribosomal:RiPP:Thiopeptide |
| Region 1.14 | NI siderophore, NRPS, transAT-PKS, T1PKS | 2,821,453 | 2,946,165 | High | kirromycin | NRPS:Type I+PKS:Type I |
| Region 1.15 | lassopeptide | 3,310,910 | 3,333,512 | High | citrulassin D | ribosomal:RiPP |
| Region 1.16 | terpene | 3,393,343 | 3,414,401 | | | |
| Region 1.17 | T1PKS | 3,420,663 | 3,512,563 | High | phoslactomycin B | PKS |
| Region 1.18 | melanin | 3,571,750 | 3,582,274 | | | |
| Region 1.19 | T1PKS, NRPS-like, PKS-like, oligosaccharide | 3,590,825 | 3,769,574 | Medium | caniferolide A/caniferolide B/caniferolide C/caniferolide D | PKS:Type I |
| Region 1.20 | terpene | 3,827,998 | 3,852,891 | Medium | isorenieratene | terpene |

| | | | | | | |
|---|---|---|---|---|---|---|
| Region 1.21 | other, NRPS, NRPS-like | 3,865,922 | 3,923,608 | High | antipain | NRPS:Type I |
| Region 1.22 | butyrolactone | 4,010,964 | 4,021,824 | | | |
| Region 1.23 | lanthipeptide-class-i | 4,118,058 | 4,150,567 | | | |
| Region 1.24 | NAPAA | 4,221,806 | 4,255,797 | High | ε-Poly-L-lysine | NRPS:Type I |
| Region 1.25 | atropopeptide | 4,455,548 | 4,476,828 | | | |
| Region 1.26 | NRPS,betalactone | 4,480,157 | 4,533,515 | | | |
| Region 1.27 | butyrolactone | 4,586,514 | 4,597,344 | | | |
| Region 1.28 | T1PKS, hglE-KS | 4,761,611 | 4,813,176 | | | |
| Region 1.29 | terpene | 4,896,728 | 4,923,457 | Medium | hopene | terpene |
| Region 1.30 | T3PKS | 5,192,667 | 5,233,848 | | | |
| Region 1.31 | T2PKS | 5,273,738 | 5,346,277 | High | lysolipin I | PKS |
| Region 1.32 | RiPP-like | 5,393,829 | 5,404,671 | | | |
| Region 1.33 | butyrolactone | 5,499,392 | 5,510,369 | | | |
| Region 1.34 | NI-siderophore | 5,818,525 | 5,851,228 | Low | kinamycin | PKS |
| Region 1.35 | triceptide | 6,375,354 | 6,396,934 | | | |
| Region 1.36 | linaridin | 7,259,696 | 7,280,606 | Medium | legonaridin | ribosomal:RiPP |
| Region 1.37 | polyhalogenated-pyrrole, NRPS, HR-T2PKS, NRPS-like, lassopeptide | 7,381,056 | 7,469,607 | Low | colibrimycin | NRPS:Type I+other:other |
| Region 1.38 | terpene | 7,470,668 | 7,492,911 | | | |
| Region 1.39 | terpene | 7,867,125 | 7,888,099 | | | |
| Region 1.40 | triceptide, lanthipeptide-class-i | 7,939,585 | 7,967,068 | | | |
| Region 2.10 | hydrogen-cyanide | 36,982 | 49,872 | Low | aborycin | ribosomal:RiPP |

**Table S6.** Annotation of biosynthetic genes of lysolipins in *Streptomyces* sp. P8-2B18.

| ORF | Size (DNA) | Length (aa) | Proposed function | ID/SI | Protein homologue and origin |
|---|---|---|---|---|---|
| *1* | 1005 | 334 | LacI family transcription regulator | 90/94 | WP_366045776.1, *Streptomyces lydicus* |
| *2* | 1539 | 512 | right-handed parallel beta-helix repeat-containing protein | 96/97 | WP_381469703.1, *Streptomyces kronopolitis* |
| *3* | 615 | 204 | PadR family transcriptional regulator | 96/98 | WP_235447681.1, Streptomyces sioyaensis |
| *4* | 1374 | 457 | FAD-dependent oxidoreductase | 92/96 | WP_392995217.1, *Streptomyces lydicus* |
| *5* | 141 | 46 | Hypothetical protein | 89/89 | WP_392995217.1, *Streptomyces kronopolitis* |
| *6* | 303 | 100 | antibiotic biosynthesis monooxygenase family protein | 91/94 | WP_235447678.1, *Streptomyces sioyaensis* |
| *7* | 360 | 119 | antibiotic biosynthesis monooxygenase family protein | 94/97 | WP_433858753.1, *Streptomyces kronopolitis* |
| *lypA* | 753 | 250 | SDR family NAD(P)-dependent oxidoreductase | 81/87 | WP_470729399.1, *Streptomyces nigrescens* |
| *8* | 462 | 153 | anthrone oxygenase family protein | 88/92 | WP_397778897.1, *Streptomyces lydicus* |
| *lypB* | 468 | 155 | cyclase/dehydrase | 77/89 | WP_190132757.1, *Streptomyces mashuensis* |
| *lypC* | 261 | 86 | acyl carrier protein | 98/97 | WP_381469679.1, *Streptomyces kronopolitis* |
| *lypD* | 1251 | 416 | ketosynthase chain-length factor | 93/95 | WP_397778903.1, *Streptomyces lydicus* |
| *lypE* | 1293 | 430 | ketosynthase | 97/98 | WP_381469673.1, *Streptomyces kronopolitis* |
| *9* | 429 | 142 | cupin domain-containing protein | 93/95 | WP_397971155.1, *Streptomyces sioyaensis* |
| *lypF* | 336 | 111 | TcmI family type II polyketide cyclase | 98/99 | WP_364875617.1, *Streptomyces kronopolitis* |
| *lypG* | 684 | 227 | SDR family NAD(P)-dependent oxidoreductase | 87/90 | WP_397778911.1, *Streptomyces lydicus* |
| *lypH* | 1158 | 385 | glycosyltransferase | 92/94 | WP_388086366.1, *Streptomyces kronopolitis* |
| *lypI* | 1842 | 613 | asparagine synthase (glutamine-hydrolyzing) | 83/88 | WP_470729389.1, *Streptomyces nigrescens* |
| *10* | 675 | 224 | TenA family protein | 88/92 | WP_397778917.1, *Streptomyces lydicus* |

| | | | | | |
|---|---|---|---|---|---|
| *lypJ* | 1209 | 402 | cytochrome P450 | 98/98 | WP_364875607.1, *Streptomyces kronopolitis* |
| *11* | 198 | 65 | ferredoxin | 92/93 | WP_388086363.1, *Streptomyces kronopolitis* |
| *lypK* | 369 | 122 | SchA/CurD-like domain-containing protein | 92/98 | WP_382854044.1, *Streptomyces sioyaensis* |
| *12* | 447 | 148 | ALF repeat-containing protein | 75/89 | WP_466085160.1, *Streptomyces albofaciens* |
| *lypL* | 846 | 281 | NmrA family NAD(P)-binding protein | 88/89 | WP_364875602.1, *Streptomyces kronopolitis* |
| *13* | 1200 | 399 | FAD-dependent monooxygenase | 93/95 | WP_388086359.1, *Streptomyces kronopolitis* |
| *14* | 1413 | 470 | MFS transporter | 95/97 | WP_006602383.1, *Streptomyces auratus* |
| *15* | 693 | 230 | TenA family protein | 93/94 | WP_382970341.1, *Streptomyces sioyaensis* |
| *lypM* | 1221 | 406 | cytochrome P450 | 92/94 | WP_235448475.1, *Streptomyces sioyaensis* |
| *lypN* | 1065 | 354 | methyltransferase | 93/95 | WP_382970336.1, *Streptomyces sioyaensis* |
| *lypO* | 1173 | 390 | cytochrome P450 | 95/96 | WP_397971106.1, *Streptomyces sioyaensis* |
| *lypP* | 1026 | 341 | methyltransferase | 95/97 | WP_397778939.1, *Streptomyces lydicus* |
| *lypQ* | 1638 | 545 | monooxygenase FAD-binding | 94/95 | WP_359363600.1, *Streptomyces sioyaensis* |
| *lypR* | 1017 | 338 | methyltransferase | 94/95 | WP_364875583.1, *Streptomyces kronopolitis* |
| *16* | 456 | 151 | anthrone oxygenase family protein | 92/94 | WP_397971096.1, Streptomyces sioyaensis |
| *lypS* | 858 | 285 | LLM class flavin-dependent oxidoreductase | 90/94 | WP_093649208.1, *Streptomyces* sp. 2314.4 |
| *lypT* | 1017 | 338 | methyltransferase | 94/96 | WP_388086348.1, *Streptomyces kronopolitis* |
| *lypU* | 882 | 293 | NAD(P)-dependent oxidoreductase | 67/80 | HEY8374604.1, Pseudonocardiaceae bacterium |
| *17* | 390 | 129 | SchA/CurD-like domain-containing protein | 90/94 | WP_364875573.1, *Streptomyces kronopolitis* |
| *lypV* | 738 | 245 | SDR family NAD(P)-dependent oxidoreductase | 93/95 | WP_235448465.1, *Streptomyces sioyaensis* |
| *lypW* | 1377 | 458 | halogenase | 72/81 | WP_359505020.1, *Streptomyces albus* |
| *18* | 1911 | 636 | transcriptional regulator, SARP family | 90/91 | WP_397778958.1, *Streptomyces lydicus* |
| *lypX* | 768 | 255 | SDR family NAD(P)-dependent oxidoreductase | 99/99 | WP_381469594.1, *Streptomyces kronopolitis* |

| | | | | | |
|---|---|---|---|---|---|
| *lypY* | 711 | 236 | class I SAM-dependent methyltransferase | 92/92 | WP_364875563.1, *Streptomyces kronopolitis* |
| *lypZ* | 1017 | 338 | methyltransferase | 93/94 | WP_411139466.1, *Streptomyces* sp. x-80 |
